# Quantifying annotation-stratified pleiotropy and co-polygenicity between complex traits

**DOI:** 10.64898/2026.06.04.730246

**Authors:** Jiayi Qu, Tianjing Zhao, Tian Lin, Ang Li, Shouye Liu, Solal Chauquet, Peter M. Visscher, Naomi R. Wray, Loic Yengo, Jian Zeng, Hao Cheng

**Affiliations:** Institute for Molecular Bioscience, The University of Queensland, Brisbane, Queensland, Australia; Big Data Institute, Li Ka Shing Centre for Health Information and Discovery, Nuffield Department of Population Health, University of Oxford, Oxford, UK; Department of Psychiatry, University of Oxford, Oxford, UK; Department of Animal Science, University of California, Davis, Davis, CA, USA; Department of Animal Science, University of Nebraska–Lincoln, Lincoln, NE, USA; Graduate Group in Biostatistics, University of California, Davis, Davis, CA, USA; Graduate Group in Integrative Genetics and Genomics, University of California, Davis, Davis, CA, USA

## Abstract

Understanding shared genetic architecture is essential to interpreting disease comorbidities and trait correlations. We introduce SBayesAPP, a Bayesian model that integrates GWAS summary statistics with functional annotations to jointly estimate annotation-stratified SNP effect-size correlation and pleiotropic variant proportion (co-polygenicity) between traits, dissecting genetic correlation and coheritability enrichment across annotations. Simulations and real data analyses show improved accuracy and interpretability over existing methods. In type 2 diabetes analyses with 15 traits, SBayesAPP reveals clear tissue- and cell-type-specific enrichment and distinguishes mechanisms driven by few large-effect variants versus many modest-effect variants. The analysis of smoking and lung cancer prioritizes lung and immune cells, and identifies cell-type-specific genetic correlations driven by either pleiotropic or lung-cancer-specific variants, consistent with a causal relationship model. For schizophrenia and educational attainment, despite near-zero genome-wide genetic correlation, cell-type-specific correlations range from −0.20 to 0.21, with strong (co)heritability enrichment and high co-polygenicity found in dopaminergic neurons and oligodendrocytes. These results highlight the ability of SBayesAPP to resolve annotation-specific genetic sharing and uncover biological mechanisms across complex traits.

## Introduction

Genome-wide association studies (GWASs) have identified thousands of associations between genetic variants and complex traits, thus revealing the polygenic architecture of these phenotypes. GWAS have shown significant heritability enrichments in functionally important genomic regions, linking GWAS associations to biological mechanisms involving trait-relevant pathways, tissues, and cell types^1^. Recent methodological advances^2–6^ have further enabled the decomposition of heritability enrichment into contributions from increased polygenicity, larger effect sizes, or both, thereby facilitating biological interpretation.

Across traits, widespread genetic correlations are observed^7^, suggesting that many genetic variants are associated with multiple, often distinct, traits^7–9^. Like genome partitioning of SNP-based heritability^10^, recent efforts^11–13,16–17^ have focused on local genetic correlations uncovering substantial variation across chromosomal segments and functional annotations, deepening our understanding of the shared genetic architecture of complex traits. However, a genetic correlation, defined as the correlation coefficient of additive genetic effects between two traits, can arise either from the same variants affecting both traits (pleiotropy), or from distinct but correlated variants in linkage disequilibrium (LD), with the latter being less biologically informative. Moreover, even when the genome-wide genetic correlation is close to zero, traits can still be genetically connected through shared causal variants with inconsistent directions of effects. Distinguishing these scenarios, pleiotropy versus LD-induced correlation or shared variants despite a null genome-wide genetic correlation, is crucial for elucidating etiological links between diseases, identifying pathways and cell types involved in both traits, informing drug repurposing, refining genetic risk profiling, and improving disease stratification.

Cross-trait linkage disequilibrium score regression (LDSC)^14^ is widely used to estimate genome-wide genetic correlation from GWAS summary statistics. High-Definition Likelihood (HDL)^15^ extends LDSC by utilizing the full LD matrix, improving the precision of genetic correlation estimates. To move beyond genome-wide averages, several methods estimate local genetic correlations (e.g., 𝜌-HESS^16^, LAVA^17^). By incorporating genomic annotations, GNOVA^11^ stratifies genetic covariance across annotations, and its applications have, for example, identified significant genetic covariance between late-onset Alzheimer’s disease and amyotrophic lateral sclerosis in predicted functional regions. However, these approaches do not distinguish whether genetic correlation arises from pleiotropy versus LD-induced correlation. MiXeR^18^ addresses this by estimating the proportions of pleiotropic and trait-specific variants via a mixture model, but it does not incorporate genomic annotations and therefore cannot localize sharing to specific functional categories. Consequently, there remains a gap for methods that can infer polygenicity and pleiotropy stratified by genomic annotations across traits.

In this study, we introduce SBayesAPP (Summary-data-based Bayesian method leveraging biological Annotations to quantify Pleiotropy and Polygenicity), which integrates genome-wide functional annotations with GWAS summary statistics while explicitly accounting for the sparsity in both trait-shared and trait-specific genetic effects. Our approach partitions genetic architecture by annotation groups to estimate annotation-stratified genetic correlation and coheritability (the ratio of genetic covariance to phenotypic covariance between two traits) enrichment. SBayesAPP further dissects these signals into contributions from polygenicity and pleiotropy by estimating, within each annotation group, both the proportion of pleiotropic variants and the correlation of effect sizes between traits. We evaluate performance through extensive simulations and comparisons with existing methods (Table 1). Using genomic annotations derived from single-cell RNA-seq data in human, we then apply SBayesAPP to the largest available GWAS summary statistics for type 2 diabetes (T2D)^19^ together with 15 traits selected based on prior T2D studies^20–42^, with the aim to identify key cell types that mediate genetic correlations with T2D through shared causal variants, strong pleiotropy, or both. Finally, we analyze lung cancer–smoking with lung-derived annotations and schizophrenia–educational attainment with brain-derived annotations to explore the major cell types that underlie these trait–trait correlations.

**Table 1.** Summary of methods used in this study and their key characteristics.

| Method | Annotation support | Model assumption | Bi-trait analysis | Parameter resolution | Heritability | Genetic correlation | Shared polygenicity<br>SNP effect correlation |
| --- | --- | --- | --- | --- | --- | --- | --- |
| LDSC | No | Infinitesimal | Yes | Genome-wide | Yes | Yes | No |
| HDL | No | Infinitesimal | Yes | Genome-wide | Yes | Yes | No |
| S-LDSC | Yes | Infinitesimal | No | Genome-wide,<br>Annotation-stratified | Yes | No | No |
| GNOVA | Yes | Infinitesimal | Yes | Annotation-stratified | Yes | Yes | No |
| SBayesAPP | Yes | Sparse mixture | Yes | Genome-wide,<br>Annotation-stratified | Yes | Yes | Yes |

## Results

### Method overview

SBayesAPP is a Bayesian mixture model that integrates functional annotations to estimate annotation-stratified genetic architecture using GWAS summary statistics and an LD correlation matrix from a reference panel (**Methods**). It models the joint effect-size distribution for two traits, allowing each SNP to have zero or nonzero effects on either trait. This formulation yields direct, data-driven estimates of the proportions of trait-1-specific (𝜋_1_), trait-2-specific (𝜋_2_), and pleiotropic (𝜋_12_) variants, together with their effect size (co)variance matrix. In the univariate setting, polygenicity is the proportion of causal variants among all variants (i.e., for trait 1: 𝜋_1_ + 𝜋_12_; for trait 2: 𝜋_2_ + 𝜋_12_). In the bivariate setting, we define co-polygenicity as 𝜋_12_, the proportion of pleiotropic variants among all variants. We also report estimated polygenicity among causal variants (i.e., polygenicity among trait-associated variants), defined as 𝜋_1_/(1 − 𝜋_0_), 𝜋_2_/(1 − 𝜋_0_), and 𝜋_12_/(1 − 𝜋_0_), where 𝜋_0_ = 1 − 𝜋_12_ − 𝜋_1_ − 𝜋_2_. To quantify the magnitude of pleiotropy among pleiotropic variants, we report the correlation of effect sizes of pleiotropic variants, estimated from their (co)variance matrix. To incorporate functional annotation, SBayesAPP models each SNP effect as a sum of annotation-specific components, allowing the joint effect-size distribution to vary across annotation groups. This framework enables estimation of annotation-stratified genetic correlations and coheritability enrichments that are fully conditional on all other annotations, thereby appropriately accounting for overlapping annotation effects.

Compared to existing cross-trait approaches, SBayesAPP introduces three key features. First, following SBayesRC^4^, a single-trait model that incorporates functional annotations, SBayesAPP employs a low-rank model to remove noise in LD, improving computational efficiency and robustness against LD differences between GWAS and reference samples. Second, SBayesAPP estimates genetic correlations and coheritability enrichments using annotation-specific joint SNP effects, accounting for both LD among SNPs and the influence of overlapping annotations. This joint modeling framework is expected to yield more accurate estimates than existing methods that are based on marginal SNP effects and LD scores. Third, in addition to estimating coheritability enrichment, SBayesAPP quantifies annotation-specific co-polygenicity and SNP effect correlation, thereby enabling the decomposition of coheritability into contributions from polygenicity, pleiotropy, and linkage, as shown in **Figure 1**. Together, these features provide deeper insights into the structure of cross-trait genetic associations and offer a more interpretable view of the genetic architecture underlying complex traits.

**Figure 1.**
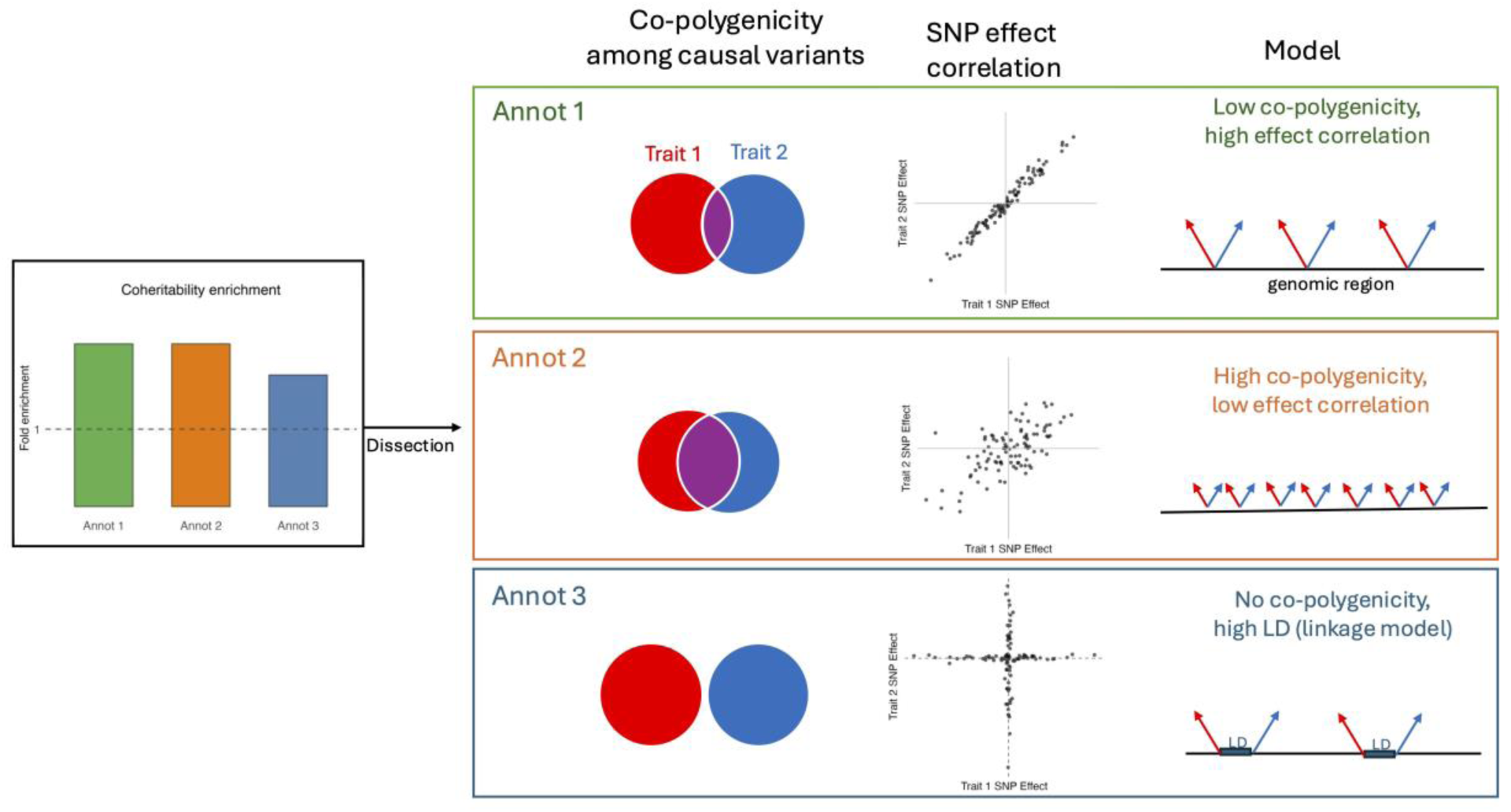
Sources of annotation-stratified coheritability enrichment: co-polygenicity, effect-size correlation, and linkage. Left: Example of estimated coheritability enrichment for three annotation groups (Annot1–Annot3), each arising from a distinct genetic architecture (schematized at right). Right: Decomposition of enrichment into two components: (i) co-polygenicity among causal variants and (ii) SNP effect correlation among pleiotropic variants, under a simplifying setting that, across annotations, the numbers of causal variants affecting Trait 1 and causal variants affecting Trait 2 are equal. The “Co-polygenicity among causal variants” column shows the proportion of pleiotropic variants (purple) among all causal variants; the “SNP effect correlation” column shows the concordance of effect sizes across pleiotropic variants. The “Model” column graphically depicts causal structure (red: effects on Trait 1; blue: effects on Trait 2) for the three genetic architectures: (i) Annot 1: low co-polygenicity, high effect-size correlation, in which coheritability enrichment is primarily driven by strong concordance among few pleiotropic variants; (ii) Annot 2: high co-polygenicity, low effect-size correlation, in which coheritability enrichment is driven by many pleiotropic variants with modest concordance in effect sizes. (iii) Annot 3 (linkage): no causal variants are shared between traits, and the apparent enrichment arises from distinct trait-specific variants in LD.

### Simulation based on real genotypes

We evaluated the performance of SBayesAPP through simulations using HapMap3 SNPs on chromosome 1 (**Methods**). We first compared the estimation accuracy of genome-wide genetic correlation (𝑟_𝑔_), genetic covariance (𝑐_𝑔_), and trait-specific heritability (ℎ_t1_^2^, ℎ^2^) among three methods: SBayesAPP, HDL, and LDSC (**Figure 2a**). Across a range of levels of co-polygenicity among causal variants (i.e., 0.1, 0.5, 0.8, and 1.0), SBayesAPP consistently produced unbiased estimates of genetic correlation and covariance, with lower estimation variance than the other methods. HDL yielded nearly unbiased genetic correlation estimates but exhibited mild overestimation when all causal variants are pleiotropic and showed greater estimation variance. HDL-estimated genetic covariance was slightly downward-biased, likely due to its underestimation of heritability. In contrast, LDSC produced significantly downward-biased estimates across all parameters and scenarios, with the highest estimation variance. For trait-specific heritability, SBayesAPP again demonstrated unbiased estimation with the highest accuracy, whereas both HDL and LDSC systematically underestimated heritability, with LDSC showing the most pronounced bias.

**Figure 2.**
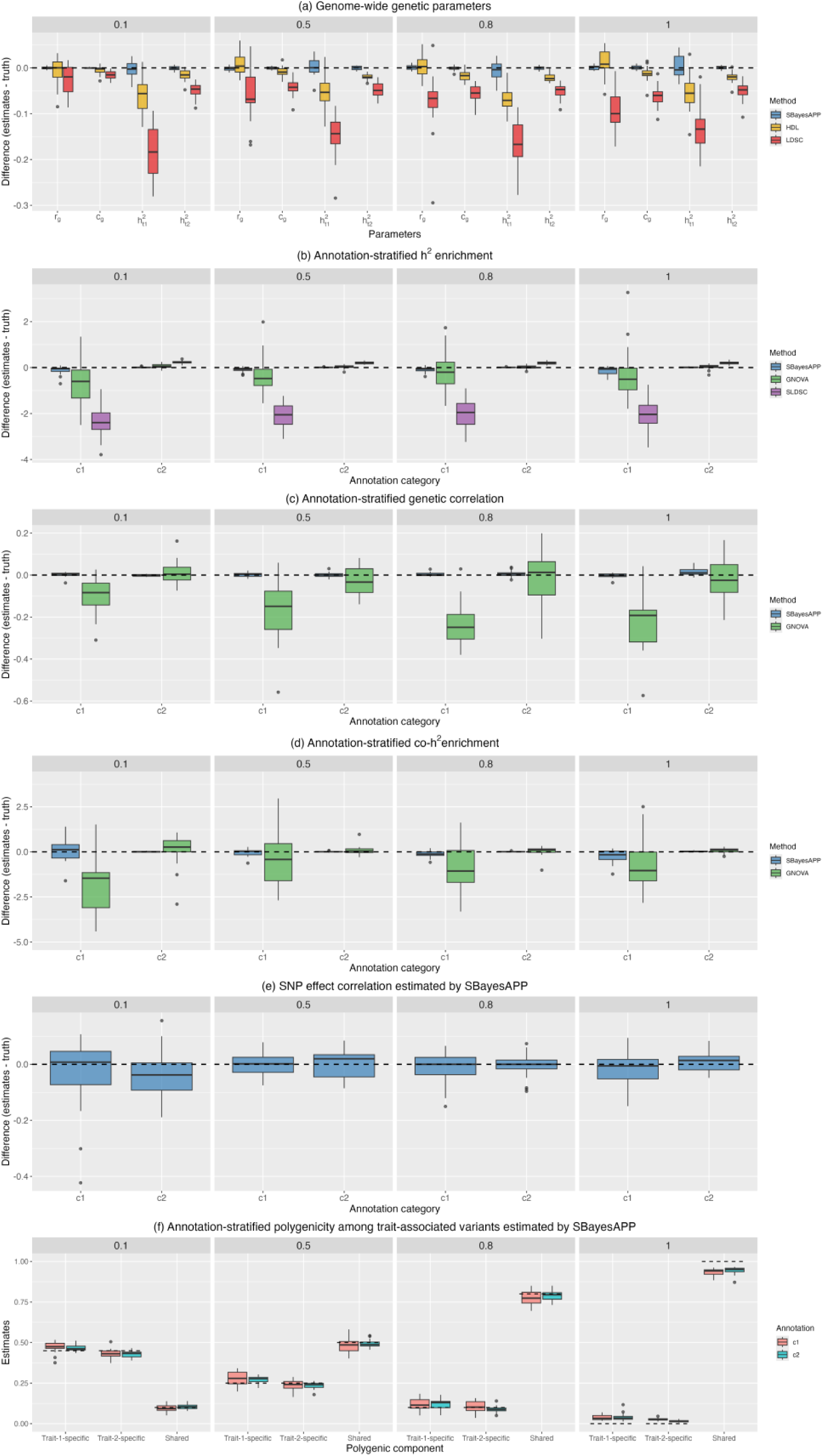
Accuracy comparison of genetic parameter estimates across methods. Panels (a–e) show the difference between estimates and true values and panel (f) shows the estimates themselves. Results are based on 20 independent simulation replicates under varying levels of co-polygenicity among causal variants (0.1, 0.5, 0.8, and 1.0). (a) Genome-wide genetic correlation (𝑟_𝑔_), genetic covariance (𝑐_𝑔_), and trait-specific heritability (ℎ_t1_^2^, ℎ^2^) from SBayesAPP, HDL, and LDSC. (b) Annotation-stratified heritability enrichment for two annotations (C1 and C2) from SBayesAPP, GNOVA, and SLDSC. Each method yields two estimates per replicate (one per trait), giving 40 data points per method per annotation and per shared-polygenicity setting. (c) Annotation-stratified genetic correlation from SBayesAPP and GNOVA. (d) Annotation-stratified coheritability enrichment from SBayesAPP and GNOVA. Thirteen extreme outliers from GNOVA (|difference| > 5), mostly from the smaller annotation (C1), were omitted for clarity. (e) Annotation-stratified SNP effect correlation from SBayesAPP. (f) Annotation-stratified polygenicity among trait-associated variants from SBayesAPP, showing the proportions of trait-1-specific, trait-2-specific, and shared variants among all trait-associated variants. Dashed lines mark the true values. Note: box-plot spread reflects variability across the 20 replicates, enabling fair comparison across methods despite differences in statistical frameworks (frequentist vs. Bayesian).

In our simulations, SNPs were partitioned into two non-overlapping annotation groups based on real annotation data (**Methods**): 9% of SNPs were assigned to C1 and 91% to C2. Within each annotation, 1% of SNPs were randomly selected as causal. To induce enrichment differences, we scaled per-SNP effects so that each annotation explained 50% of the total heritability for both traits, regardless of its size. Among pleiotropic variants, the effect-size correlation was set to 0.8 in C1 and 0.2 in C2. Thus, C1 represents a “low co-polygenicity, high effect-size correlation” scenario and C2 a “high co-polygenicity, low effect-size correlation” scenario (**Figure 1**).

Comparing differences between estimated and true values of annotation-stratified heritability enrichment across C1 and C2 (**Figure 2b**), SBayesAPP showed consistently lower bias and variation than GNOVA and S-LDSC. Next, we compared SBayesAPP and GNOVA for estimating annotation-stratified genetic correlation and coheritability enrichment (**Figure 2c-d)**. Across both annotations, SBayesAPP produced unbiased estimates and exhibited consistently lower estimation variance. GNOVA performed well for the large category (C2) but showed downward bias and reduced precision for the smaller category (C1). Notably, GNOVA produced 13 (8.1% of simulations) extreme outliers in coheritability enrichment (i.e., |estimate - truth| > 5), 12 of which occurred in C1.

By jointly estimating the annotation-stratified effect-size correlation and the mixture proportions of pleiotropic versus trait-specific components among trait-associated variants, SBayesAPP disentangles whether coheritability enrichment arises from stronger pleiotropic effect concordance, a higher proportion of shared causal variants, or both. In our simulations, both annotations contributed equally to total coheritability but differed in architectures: C1 had fewer pleiotropic variants with stronger effect-size correlation, whereas C2 had more pleiotropic variants with weaker correlation (**Methods**). SBayesAPP accurately recovered the annotation-stratified effect-size correlations across all levels of co-polygenicity among causal variants (**Figure 2e**) and yielded nearly unbiased estimates of these levels, with only a slight downward bias in the extreme case where all causal variants are pleiotropic (**Figure 2f**).

Overall, these results demonstrated the robustness of SBayesAPP for genome-wide and annotation-stratified genetic parameter estimation and its unique ability to quantify both effect-size correlation and the composition of causal variants, thereby offering valuable insights into the genetic architecture underlying cross-trait associations.

### Proof-of-principle using real data for type 2 diabetes, fasting glucose, and height

To illustrate the ability of SBayesAPP to distinguish different sources of coheritability enrichment, we performed a proof-of-principle analysis using SNP annotations defined by GWAS significance for T2D–fasting glucose and, as a biologically contrasting pair, T2D–height. SNPs were partitioned into three mutually exclusive sets: (1) significant SNP set, comprising SNPs with p-values ≤ 5 × 10^−5^ in both traits; (2) control SNP set, matched in size to the significant SNP set, and randomly sampled from SNPs with p-values > 0.5 in both traits; and (3) rest SNP set, comprising all remaining SNPs. Using SBayesAPP, we estimated, for each set, annotation-stratified coheritability enrichment, SNP effect-size correlation, co-polygenicity, and polygenicity among trait-associated variants. Because these annotations were constructed from GWAS significance, we do not aim to make new biological discoveries; rather, the goal in this section is to validate that SBayesAPP would produce expected results of enrichment in the significant SNP set (positive control group) and depletion in the control SNP set (negative control group).

**Figure 3** summarizes the results. In the T2D–fasting glucose analysis, the significant SNP set showed a 702–fold coheritability enrichment, far exceeding the 113–fold enrichment for the corresponding set in T2D–height. For both trait pairs, the rest and control SNP sets were depleted, with the rest SNP set consistently showing slightly higher fold enrichment than the control SNP set. This enriched-vs-depleted contrast is expected by the experimental design and the mechanistic differences between T2D–fasting glucose and T2D–height are shown by the effect-size correlations and variant-component distributions. For T2D–fasting glucose (r_g_ = 0.43), the effect-size correlation was strong in the significant SNP set (0.87), weaker in the rest SNP set (0.46), and insignificantly different from zero in the control SNP set. For T2D–height (r_g_ = −0.04), correlations were weaker and negative (significant: −0.23; rest: −0.09), with the control SNP set again indistinguishable from zero. In both analyses, control-SNP-set estimates of SNP effect correlation had large posterior standard deviations, indicating greater uncertainty and likely reflecting limited genetic signal in that set.

**Figure 3.**
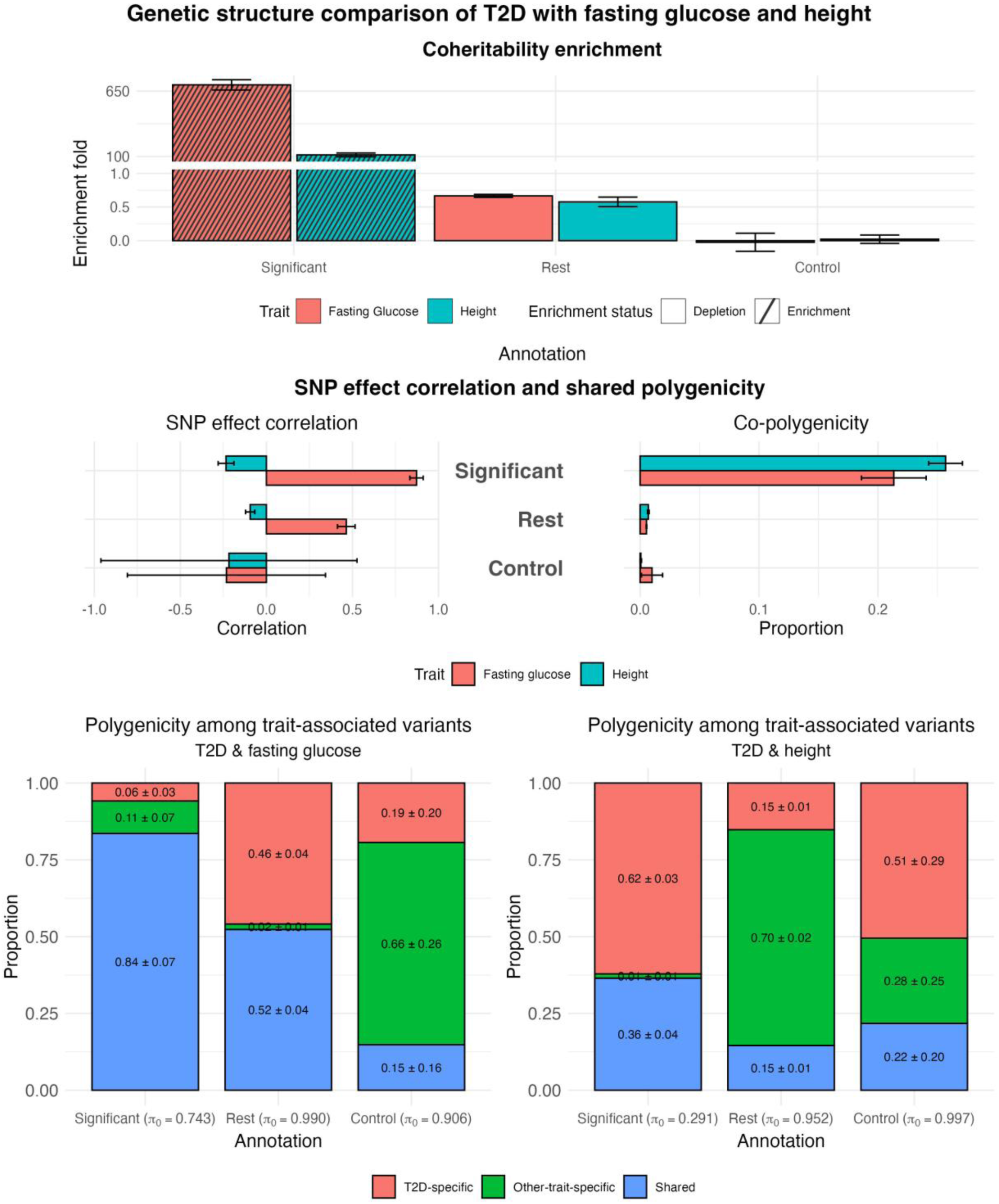
Genetic structure comparison of T2D with fasting glucose and height. This figure presents annotation-stratified analyses by SBayesAPP for T2D–fasting glucose and T2D–height based on three SNP groups: (1) Significant: SNPs with p-values ≤ 5 × 10^−5^ in both traits; (2) Control: a size-matched set of SNPs randomly sampled from those with p-values > 0.5 in both traits; and (3) Rest: all remaining SNPs not in the significant or control groups. The top row shows coheritability fold enrichment estimates, with error bars indicating posterior standard deviations from MCMC samples. Slash-filled bars indicate annotations with enriched signals; solid bars indicate depletion. The middle row presents a decomposition of these enrichments into SNP effect correlation and co-polygenicity. Red bars correspond to the T2D–fasting glucose results, and teal bars to the T2D–height results. The bottom row displays the polygenic composition among trait-associated variants within each annotation, partitioned into trait-specific and shared (pleiotropic) components. π₀ values (the estimated proportion of null SNPs) are shown beneath each annotation. Together, these results reveal distinct patterns of shared genetic architecture between the two trait pairs.

Co-polygenicity (𝜋_12_) in the significant SNP set was 0.21 for T2D–fasting glucose and 0.26 for T2D–height. In both analyses, the proportion of null SNPs (𝜋_0_) was lower in the significant SNP set than in the control SNP set, and the share of trait-associated variants (1-𝜋_0_) in the significant SNP set was 25.7% in T2D–fasting glucose and 70.9% in T2D–height. Conditional on being trait-associated, the polygenicity among trait-associated variants revealed divergent architectures: in T2D–fasting glucose, 84% of trait-associated variants in the significant SNP set were pleiotropic, whereas in T2D–height, only 36% were pleiotropic, with 62% being T2D-specific, indicating a more trait-specific architecture. In the control SNP sets, large posterior standard deviations for co-polygenicity among trait-associated variants indicated weak shared signal. In the rest SNP set, 52% of trait-associated variants were shared in T2D–fasting glucose (consistent with its stronger genome-wide correlation), whereas 70% were height-specific for T2D–height (consistent with the high polygenicity of height). Finally, because the height GWAS sample size is ∼10 × larger than that for fasting glucose (**Supplementary Table 1**), more trait-associated (both trait-specific and shared) variants are expected to be detected in the T2D–height analysis.

### Cell-type analysis in type 2 diabetes and fasting glucose

Next, we investigated the shared genetic architecture between T2D and fasting glucose using 155 cell type annotations derived from a human single-cell RNA-seq dataset^43^. For interpretability, we grouped cell types into seven tissue categories, including six T2D-relevant categories: adipose^44^, blood^45–48^, digestive^49–52^, immune^53–59^, liver^60^, pancreas^61–67^, and one category representing other tissues. Annotations in the “other” category are considered to represent tissues of secondary relevance to the shared genetic basis of T2D and fasting glucose based on the literature.

We compared coheritability enrichment estimates between SBayesAPP and GNOVA for the T2D–fasting glucose pair (**Figure 4a-b**). For SBayesAPP, uncertainty was quantified using posterior standard deviations derived from 3,000 MCMC iterations across 10 independent chains. GNOVA estimates were accompanied by standard errors approximated via the delta method, based on the standard errors of corrected genetic covariance.

**Figure 4.**
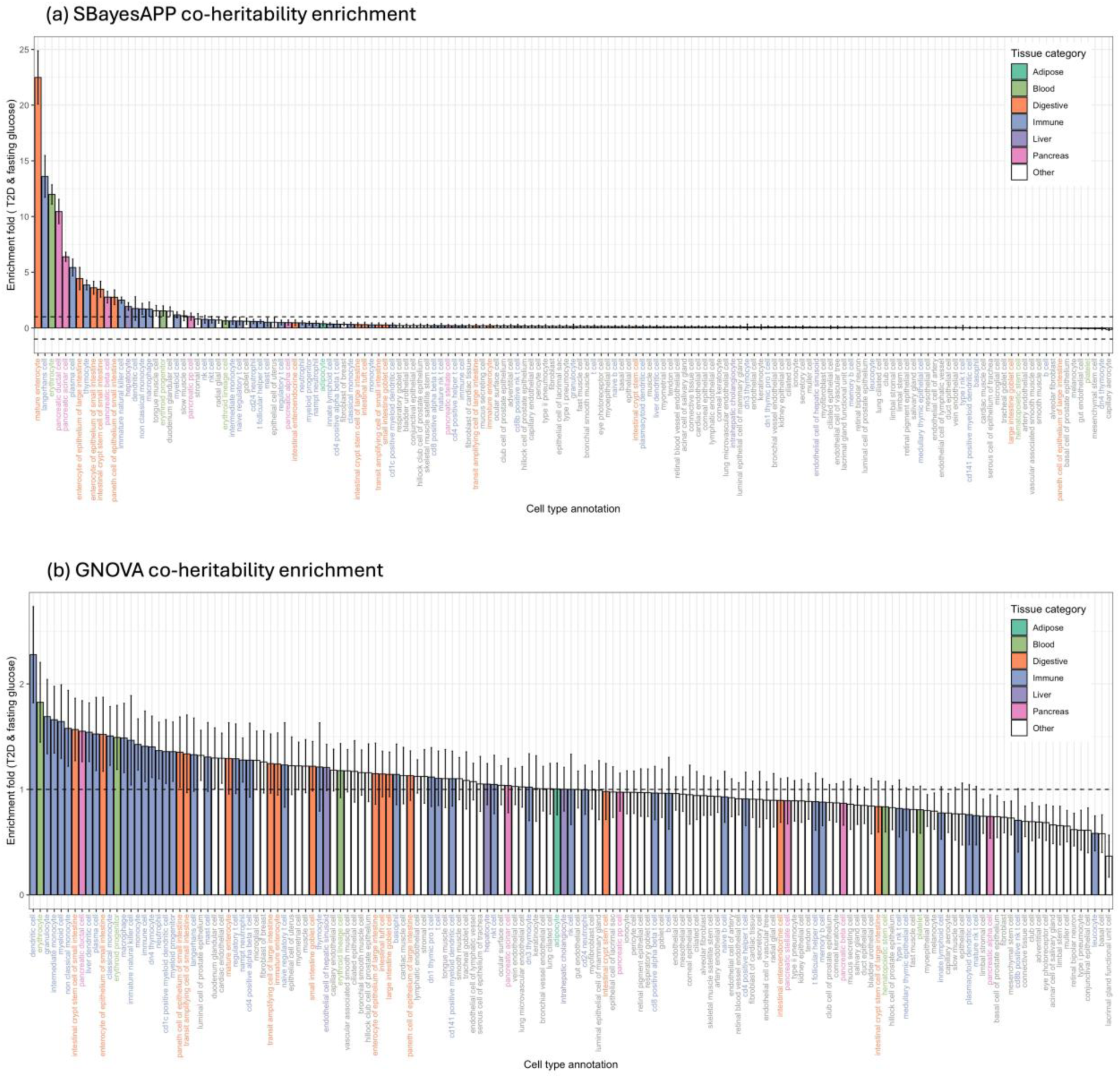
Dissecting coheritability enrichment across cell type annotations for T2D and fasting glucose. (a-b) Annotation-stratified coheritability enrichment estimates from SBayesAPP (a) and GNOVA (b) for the trait pair T2D and fasting glucose. SBayesAPP estimates represent posterior means with posterior standard deviations computed from 3,000 MCMC iterations across 10 independent chains. GNOVA estimates are shown with standard errors approximated using the delta method. (c) Decomposition of coheritability enrichment by SBayesAPP for annotations identified as significantly enriched (posterior probability > 0.95 that the fold enrichment exceeds 1). The left panel shows estimated coheritability enrichment; the middle panel shows estimated SNP effect correlation; and the right panel shows estimated co-polygenicity. Error bars denote posterior standard deviations. (d) Polygenicity among trait-associated variants for annotations that are enriched in T2D–fasting glucose analysis. Stacked bars show the contributions of shared (blue), T2D-specific (red), and other-trait-specific (green) SNPs to the total genetic signal within each annotation. The left panel shows estimates from the T2D–fasting glucose analysis; the right panel shows corresponding estimates for the same set of annotations from the T2D–height analysis. Note that the annotations shown are selected based on enrichment in T2D–fasting glucose only. All annotations are color-coded by tissue category (Adipose, Blood, Digestive, Immune, Liver, Pancreas, and Other) in all panels.

Among the top signals, three pancreatic cell types, beta cells, acinar cells, and ductal cells, were identified by SBayesAPP as significantly enriched for coheritability between T2D and fasting glucose. Significance was defined as having a posterior probability > 0.95 that the fold enrichment exceeds 1. These findings are consistent with known biological roles: pancreatic beta cells are central to glucose homeostasis through insulin secretion^65–67^; acinar cells influence broader aspects of systemic metabolic regulation by interacting with beta cells^63–64^; and ductal cells have been implicated in glucose dysregulation, particularly under pathological conditions such as T2D^61–62^. In contrast, GNOVA identified significant enrichment only in pancreatic ductal cells. In GNOVA, significance was determined based on whether the lower bound of the 90% confidence interval exceeds one. While both SBayesAPP and GNOVA prioritized cell types from multiple T2D-relevant tissues, the magnitude of fold enrichment varied considerably. GNOVA estimates typically clustered around 1, making it challenging to distinguish clearly between enriched and depleted annotations. In contrast, SBayesAPP provided more distinct separation among these annotations.

Beyond enrichment, SBayesAPP enables deeper dissection of genetic sharing by estimating annotation-specific SNP effect correlations and co-polygenicity. While existing methods often conflate genetic correlation with pleiotropic effect-size correlation, SBayesAPP decomposes the shared signal to determine whether enrichment is driven by a few highly correlated variants or many modestly correlated variants. **Figure 4c** presents this decomposition for significantly enriched annotations identified by SBayesAPP, displaying coheritability enrichment, SNP effect correlation, and co-polygenicity. Notably, the patterns for these three metrics varied substantially between annotations, reflecting differences in underlying genetic architecture.

For instance, mature enterocytes showed the strongest coheritability enrichment, the highest SNP effect correlation, and modest co-polygenicity. Enterocytes are essential for nutrient absorption and glucose transport via GLUT2, and dysregulation of insulin-mediated GLUT2 expression has been linked to hyperglycemia and metabolic dysfunction^68–69^. Lipid processing alterations and chylomicron overproduction in enterocytes further contribute to the pathophysiology of T2D^70^.

In contrast, erythrocytes ranked third in coheritability enrichment but exhibited the highest co-polygenicity and lower SNP effect correlation than mature enterocytes. Although erythrocytes are not directly involved in glucose regulation, changes in their structure and function, such as altered deformability and aggregation, are known consequences of metabolic abnormalities in T2D^71–72^. The high co-polygenicity observed in this annotation likely reflects a broad distribution of variants influencing erythrocyte-related traits in the context of systemic metabolic regulation.

In addition, we reported polygenicity among trait-associated variants. The left panel of **Figure 4d** illustrates the relative proportions of shared, T2D-specific, and fasting glucose-specific (i.e., other-trait-specific) variants among trait-associated variants within each enriched annotation.

Many of these annotations showed high levels of co-polygenicity among trait-associated variants, with some reaching 58%, consistent with the strong genome-wide genetic correlation between T2D and fasting glucose. To contrast this with a more genetically distinct trait pair, we examined the polygenicity among trait-associated variants for the same set of annotations in a T2D–height analysis. As shown in the right panel of **Figure 4d**, these annotations revealed predominantly trait-specific architectures. In particular, T2D-specific variants contributed most of the polygenic signal among the pancreatic cell types, where pancreatic beta cells (81%) and pancreatic ductal cells (78%) showed the highest proportions of T2D-specific associated variants.

Together, our results demonstrated that SBayesAPP not only detects coheritability enrichment but also distinguishes whether such enrichment arises from polygenic sharing or effect-size correlation. This capability enables nuanced interpretation of pleiotropic mechanisms and offers unique insights into the genetic basis of complex trait relationships.

#### Cell-type analysis in T2D-related traits

Beyond fasting glucose, we analyzed annotation-stratified coheritability enrichment between T2D and 14 additional traits, yielding 15 T2D–trait pairs. Traits were selected a priori from the T2D literature and grouped as follows: (i) heart-related traits: atrial fibrillation^20^ and systolic blood pressure^21^; (ii) brain-related traits: insomnia^22–23^, Parkinson’s disease^24–25^, depression^26–27^, schizophrenia^28–29^, and educational attainment^30–31^; (iii) cancer-related trait: prostate cancer^32–33^; (iv) obesity-related trait: body mass index^34–35^; (v) autoimmune-related traits: rheumatoid arthritis^36–37^ and inflammatory bowel disease^38–39^; and (vi) metabolism-related trait: cholesterol^40–41^. As controls, we included height and red blood cell (negative controls) and fasting glucose (positive control). Most of these traits overlap with those examined in a recent study of T2D-related traits^42^ (**Supplementary Table 1**).

We first compared genome-wide estimates of SNP-based heritability, genetic covariance, and genetic correlation from SBayesAPP, HDL, and LDSC across all 15 T2D–trait pairs (**Supplementary Figure 1**). In most cases, HDL’s estimates were between those of SBayesAPP and LDSC, consistent with patterns observed in our simulation studies. SBayesAPP generally yielded higher SNP-based heritability estimates than both LDSC and HDL, with exceptions for fasting glucose, cholesterol, and Parkinson’s disease. For genetic covariance, LDSC typically reported the largest magnitude (by absolute value), followed by HDL and then SBayesAPP.

**Figure 5** shows SBayesAPP-estimated coheritability enrichment for T2D with 15 secondary traits. Traits on the y-axis were ordered by decreasing genome-wide genetic correlation with T2D, from the highest (body mass index) to the most negative (educational attainment). Cell types on the x-axis were grouped into 18 tissue categories; six tissues of a priori relevance to T2D are outlined in black boxes. Color encodes the sign of genetic covariance and the magnitude of the enrichment z-score (defined as posterior mean divided by posterior standard deviation), and circle size encodes the co-polygenicity among trait-associated variants. SBayesAPP revealed clearer and more biologically plausible results than other methods: most enrichment concentrated in the six highlighted tissues plus muscle^73^ tissue, whereas cell types from other tissues were largely depleted. Moreover, trait pairs with stronger genome-wide genetic correlations tended to show darker colors and larger circles within enriched annotations, indicating greater coheritability enrichment and higher co-polygenicity among trait-associated variants. In addition, for strongly correlated pairs (either positive or negative), the sign of covariance in enriched annotations generally aligned with the sign of the genome-wide genetic correlation, whereas weakly correlated pairs showed a mix of positive and negative signals. These patterns indicated that genome-wide genetic correlation alone may not fully capture the complexity of shared genetic architecture across annotations. In contrast, GNOVA exhibited a more diffuse pattern with scattered signals across cell types, lacking estimates of co-polygenicity, offers less biological interpretability (**Supplementary Figure 2**).

**Figure 5.**
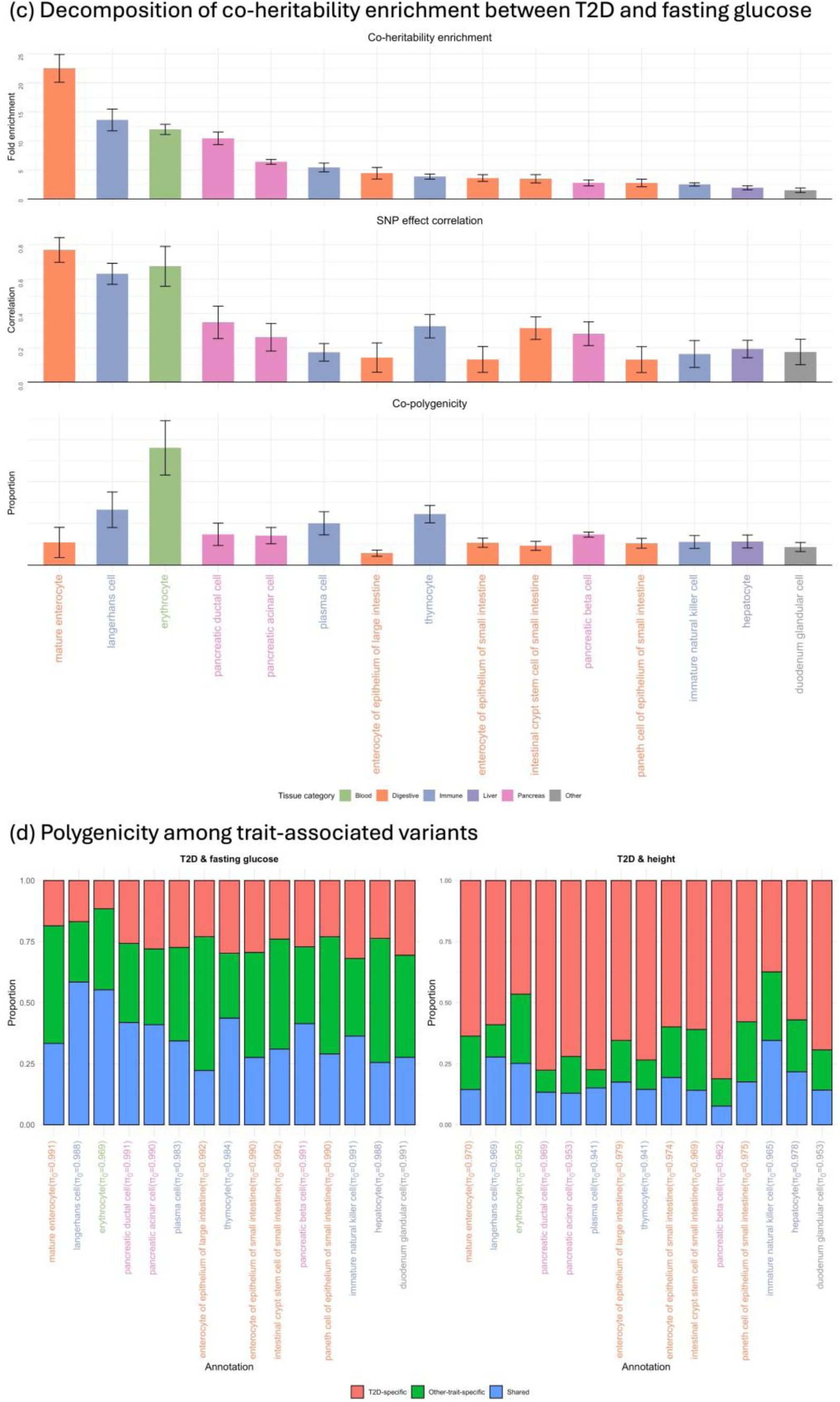
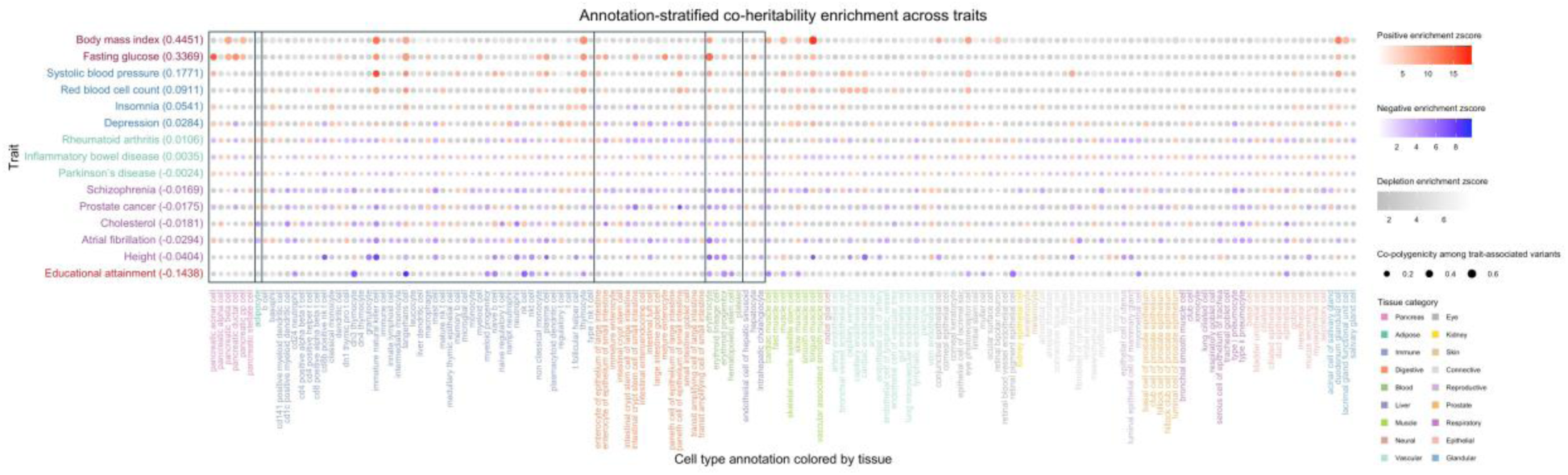
Annotation-stratified coheritability enrichment between T2D and 15 traits. This figure shows SBayesAPP-estimated coheritability enrichment for T2D with 15 secondary traits. Traits are listed on the y-axis, ordered by their genome-wide genetic correlation with T2D (shown in parentheses), from most positive (top) to most negative (bottom). The x-axis displays 155 cell type annotations grouped into 18 tissue categories, arranged in the following order: pancreas, adipose, immune, digestive, blood, liver, muscle, vascular, eye, kidney, skin, connective, reproductive, prostate, respiratory, epithelial, and glandular. The first six tissue categories of a priori relevance to T2D, are highlighted with black boxes. Each circle represents the result for a trait-annotation pair. Circle size indicates co-polygenicity among trait-associated variants, defined as the proportion of pleiotropic variants among all trait-associated variants. Larger circles correspond to higher co-polygenicity among trait-associated variants. Color indicates the sign of genetic covariance and the magnitude of coheritability enrichment z-score, defined as the posterior mean divided by the posterior standard deviation. Both red and blue represent annotations with fold enrichment greater than 1, where red corresponds to positive genetic covariance and blue to negative covariance. Darker shades of red or blue indicate higher enrichment z-scores. Gray indicates annotations with fold enrichment between –1 and 1 (i.e., depleted annotations), with darker gray representing values closer to zero (i.e., smaller z-score).

#### Analysis of smoking and lung cancer

In our subsequent real data analysis, we analyzed smoking^75^ (cigarettes per day) and lung cancer^74^ using 43 annotation groups: 42 lung-derived cell types and one background group containing all SNPs not assigned to any cell type. We selected this trait pair because the causal direction is well established, smoking increases lung cancer risk, making it a biologically interpretable test case for cross-trait analysis. We classified annotations into three categories: (i) lung-specific, upper airway and lung-resident cell types; (ii) immune, cell types broadly distributed across multiple tissues (including lung); and (iii) other, broadly distributed non-immune cell types plus the background group.

We compared results from SBayesAPP and GNOVA. Using SBayesAPP, we identified seven significantly enriched cell types (posterior probability of enrichment > 0.95; **Figure 6**): four lung-specific (epithelial cell of the lower respiratory tract, type I pneumocyte, epithelial cell of the alveolus of lung, and lung pericyte), two immune (T cell, natural killer [NK] cell), and one in the “other” category (hematopoietic stem cell). In contrast, GNOVA detected no significant enrichment at the 90% confidence interval (**Supplementary Figure 3**). Under a more liberal heuristic (enrichment estimate – 1 standard error > 1), GNOVA highlighted four cell types: three immune (T cell, plasmacytoid dendritic cell, and NK cell) and one “other” (hematopoietic stem cell). Three of these overlapped with the SBayesAPP results (T cell, NK cell, hematopoietic stem cell), underscoring the contribution of immune-related processes to the shared genetic architecture between smoking and lung cancer.

**Figure 6.**
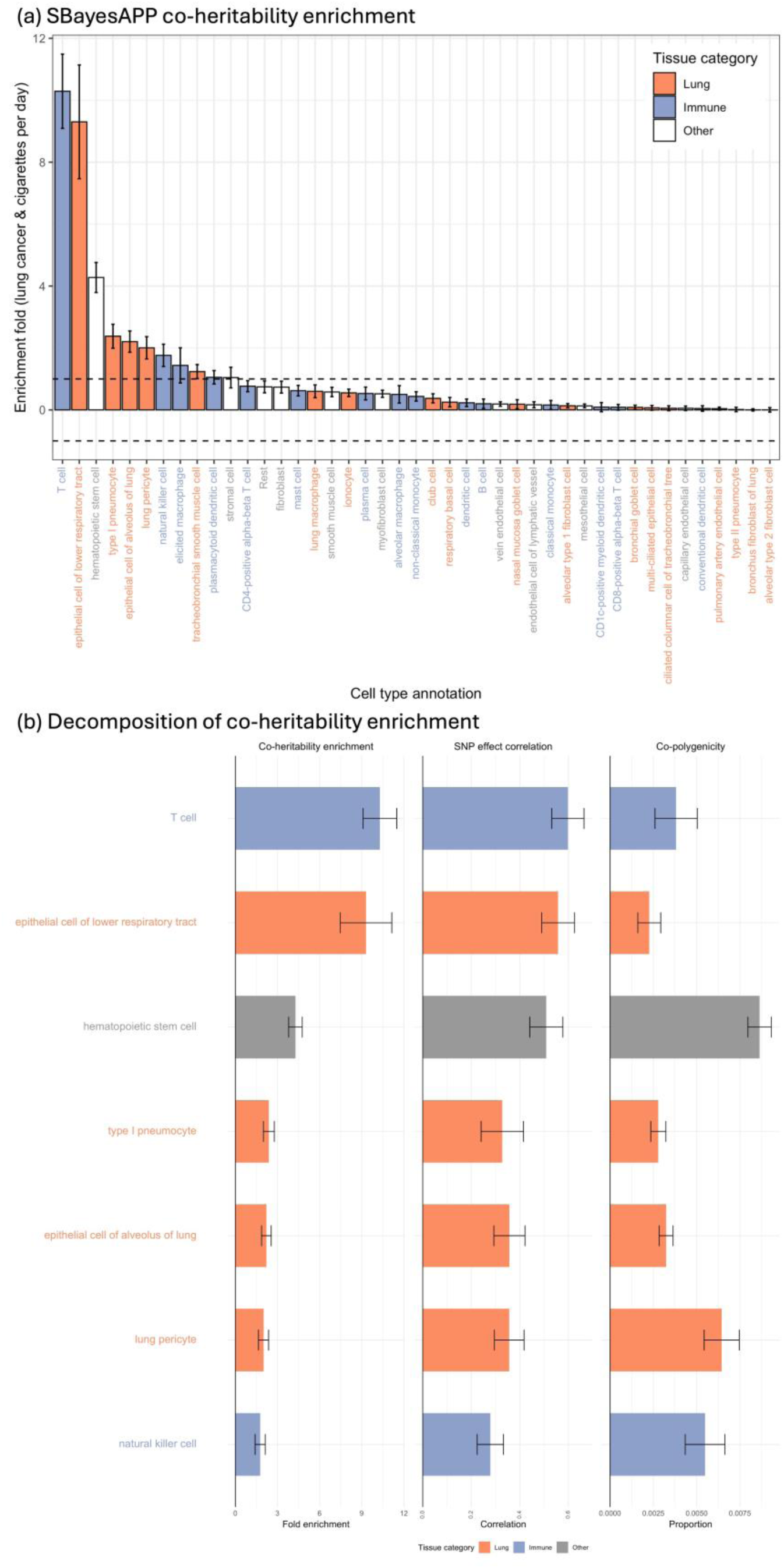
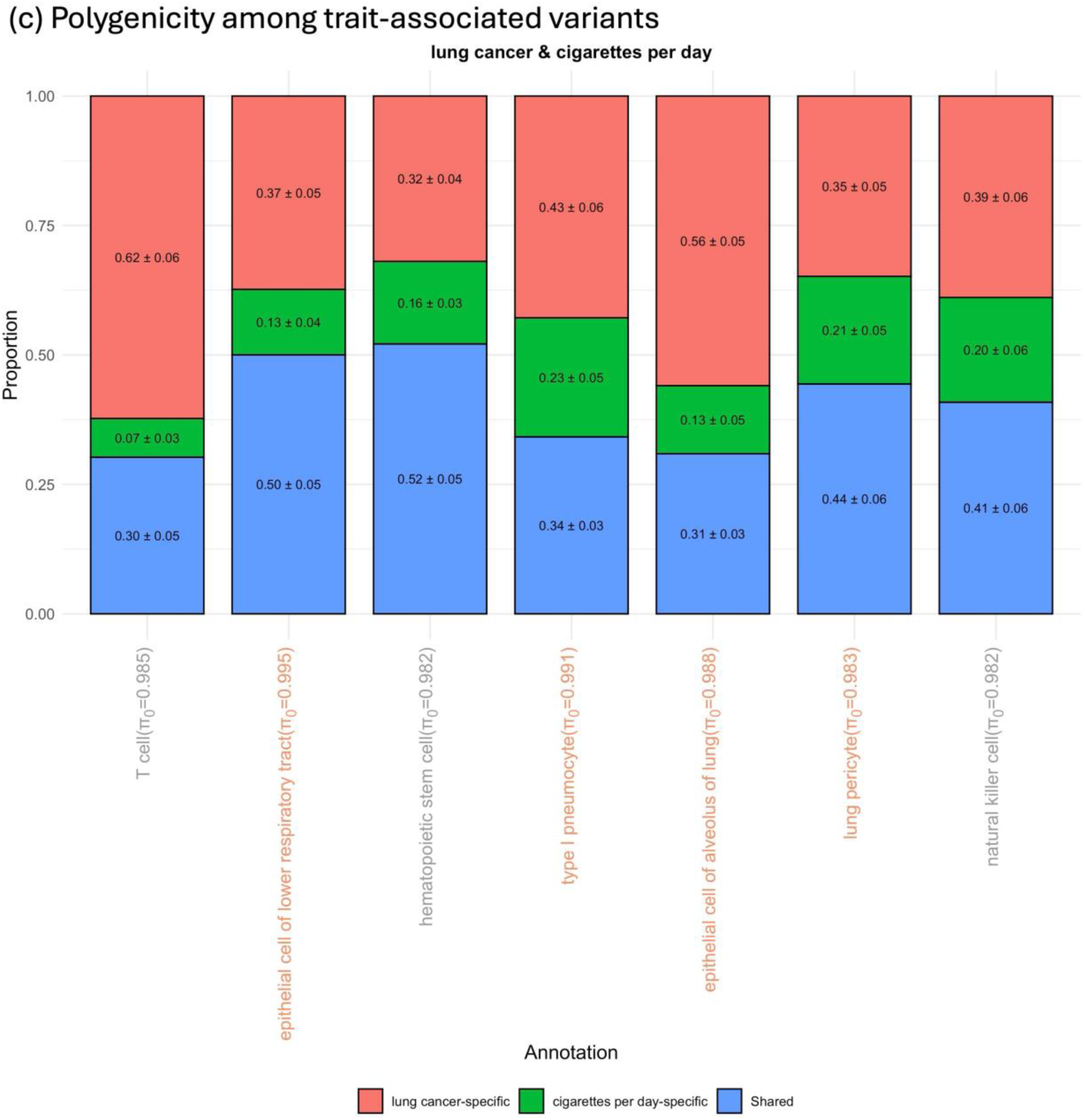
Dissecting coheritability enrichment for lung cancer and cigarettes per day. (a) Annotation-stratified coheritability enrichment estimates from SBayesAPP for the trait pair lung cancer and cigarettes per day. SBayesAPP estimates represent posterior means with posterior standard deviations computed from 3,000 MCMC iterations across 10 independent chains. (b) Decomposition of coheritability enrichment by SBayesAPP for annotations identified as significantly enriched (posterior probability > 0.95 that the fold enrichment exceeds 1). The left panel shows estimated coheritability enrichment; the middle panel shows estimated SNP effect correlation; and the right panel shows estimated co-polygenicity. Error bars denote posterior standard deviations. (c) Polygenicity among trait-associated variants for annotations identified as significantly enriched. Stacked bars show the contributions of shared (blue), lung cancer-specific (red), and cigarettes per day-specific (green) SNPs to the total genetic signal within each annotation. The estimate and its standard error are shown inside each bar. All annotations are color-coded by tissue category in all panels.

The SBayesAPP findings point to two major biological insights. First, immune-related but non–lung-specific cell types play an important role. T cells and NK cells are central to immune surveillance and tumor immunity in lung cancer^76–77^, and smoking is known to disrupt their function and promote chronic inflammation, thereby facilitating tumor initiation^78^. Smoking also affects hematopoietic stem and progenitor cells (HSPCs) by inducing oxidative stress and inflammation, leading to dysregulated hematopoiesis and impaired immune function^79–80^.

Second, several lung-specific cell types also showed significant enrichment. For example, lower-airway epithelial cells are directly exposed to cigarette smoke and display persistent gene-expression changes, even after smoking cessation, which may explain the long-lasting elevated risk of lung cancer in former smoker^81^. Likewise, alveolar epithelial cells are essential for maintaining normal alveolar structure and function; their injury from cigarette smoke toxins increases epithelial permeability, reduces surfactant production, triggers abnormal cytokine and growth factor signaling, and elevates lung cancer risk^82^.

Beyond detecting enrichment, SBayesAPP further decomposed the coheritability enrichment into annotation-specific SNP effect correlation, co-polygenicity, and polygenicity among trait-associated variants. T cells exhibited the largest SNP effect correlation, while hematopoietic stem cells showed the largest co-polygenicity. Among the trait-associated variants of the enriched annotations, most were either pleiotropic or lung-cancer-specific, with smoking-specific associated variants contributing the least. This pattern is generally consistent with the known causal role of smoking in lung cancer, where the shared genetic architecture is more strongly driven by lung cancer risk alleles and pleiotropic variants than by variants associated exclusively with smoking.

#### Analysis of schizophrenia and educational attainment

Last, we analyzed schizophrenia and educational attainment (EA) using nine annotation groups, eight brain-derived cell types plus one background group. Although the genome-wide genetic correlation between schizophrenia and EA was close to zero (r_g_ = 0.017), several cell types showed significant annotation-level genetic correlations: excitatory neurons had the strongest positive value (0.21), whereas endothelial cells had the strongest negative value (−0.20) (**Figure 7a**). In line with these directions, several cell types also showed significant coheritability enrichment: oligodendrocytes, excitatory neurons, and dopaminergic (DA) neurons were positively enriched, while microglia and endothelial cells were negatively enriched (posterior probability of enrichment > 0.95). For comparison, GNOVA detected significant coheritability enrichment only in DA neurons (**Supplementary Figure 4**). Importantly, single-trait heritability enrichment and coheritability enrichment are not interchangeable, and different cell types exhibited distinct patterns. Inhibitory neurons were not significantly enriched for coheritability, but did show significant SNP-based heritability enrichment for both traits. Conversely, excitatory neurons, microglia, and endothelial cells were enriched for coheritability but were not enriched for single-trait heritability. Oligodendrocytes and DA neurons stood out by showing enrichment for both SNP-based heritability and coheritability.

**Figure 7.**
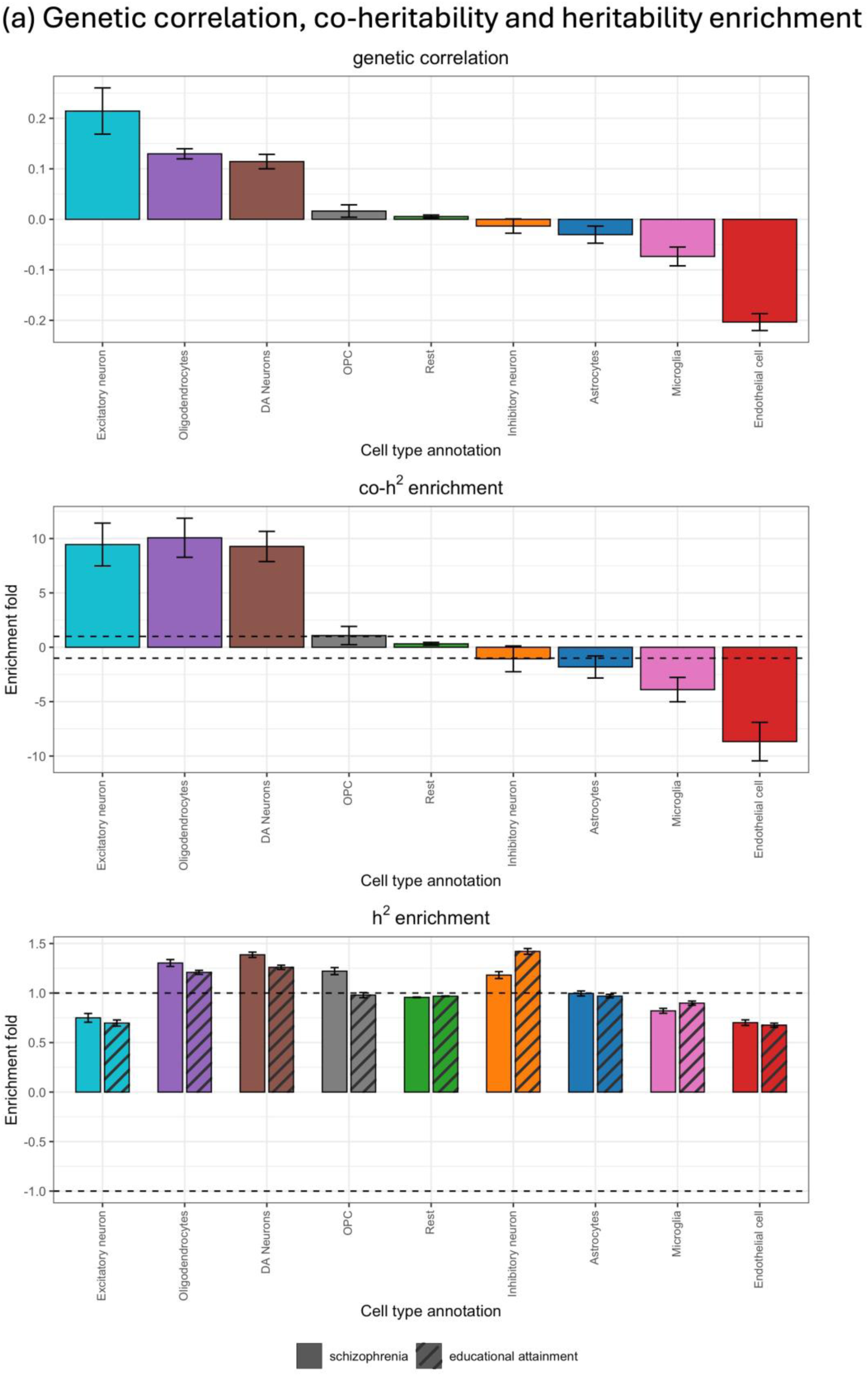
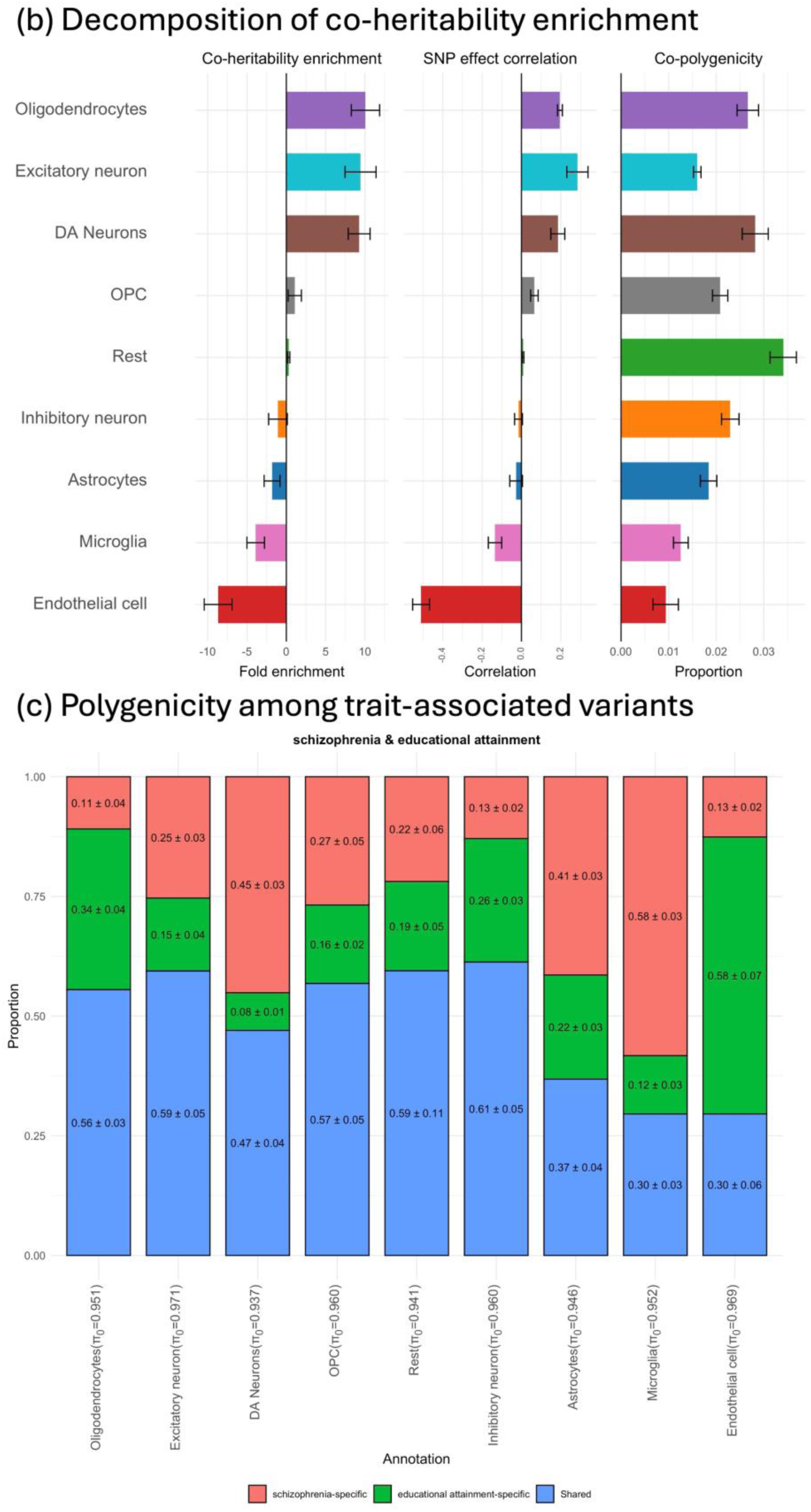
Dissecting coheritability enrichment across cell type annotations for schizophrenia and educational attainment. (a) Annotation-stratified genetic correlation, coheritability enrichment, and heritability enrichment estimated by SBayesAPP. SBayesAPP results represent posterior means with posterior standard deviations, obtained from 3,000 MCMC iterations across 10 independent chains. (b) Decomposition of coheritability enrichment by SBayesAPP. The left panel shows estimated coheritability enrichment; the middle panel shows estimated SNP effect correlation; and the right panel shows estimated co-polygenicity. Error bars denote posterior standard deviation. (c) Polygenicity among trait-associated variants. Stacked bars show the contributions of shared (blue), schizophrenia-specific (red), and educational attainment-specific (green) SNPs to the total genetic signal within each annotation. The estimate and its standard error are shown inside each bar.

Among the positively enriched annotations, excitatory neurons showed the strongest SNP effect-size correlation, while DA neurons displayed the largest co-polygenicity (**Figure 7b**). The polygenic composition among trait-associated variants varied across annotations: most groups were near 50% co-polygenicity, with microglia and endothelial cells showing the lowest levels (∼30%) (**Figure 7c**). This relatively high degree of shared trait-associated variants between schizophrenia and EA aligns with Frei et al.^18^, who reported that large numbers of shared causal variants exist between schizophrenia and EA despite minimal genome-wide genetic correlation. Microglia and endothelial cells appeared to contribute negatively to the genetic covariance; however, their polygenicity among trait-associated variants was dominated by trait-specific components: microglia were composed mainly of schizophrenia-specific variants, and endothelial cells were dominated by EA-specific variants. Taken together, these results illustrate the complexity of schizophrenia–EA genetic sharing mechanisms, with annotation-specific differences in both effect-size correlation and polygenicity.

## Discussion

In this study, we present SBayesAPP, a Bayesian framework that jointly estimates, for each annotation and conditional on all others, (i) the genetic covariance matrix (yielding genetic correlations and coheritability enrichments), (ii) the SNP effect-size covariance matrix (yielding SNP effect correlation) and (iii) the polygenic mixture proportions (yielding co-polygenicity and trait-specific polygenicity among trait-associated variants). This separation of effect-size correlation and co-polygenicity provides deeper insight into shared genetic architecture. An annotation can exhibit high coheritability enrichment either because pleiotropic effect sizes are strongly correlated (effect-size-driven enrichment) or because many variants are shared across traits despite modest effect-size concordance (polygenicity-driven enrichment). Together, these estimates enable a nuanced dissection of the sources of coheritability enrichment across functional annotations.

Among SNPs nominally significant in both single-trait GWAS, we observed contrasting patterns of coheritability enrichment for T2D–fasting glucose and T2D–height. Beyond the markedly higher enrichment in the T2D-fasting glucose pair, its signal was driven by both strong SNP effect-size correlation and high co-polygenicity among trait-associated variants. In contrast, theT2D–height pair showed weaker effect-size correlation and lower co-polygenicity among trait-associated variants. Interpretation should also consider relative GWAS power: height was chosen as a biologically contrasting trait, but its GWAS has ∼10× the sample size of Fasting glucose and ∼70× more independently associated SNPs (**Supplementary Table 1**), yielding more SNPs shown associated to both traits. Despite this greater power for height, the T2D–fasting glucose pair still exhibited higher fold enrichment, stronger SNP effect correlation, and greater co-polygenicity among trait-associated variants, indicating a more closely shared genetic architecture.

Compared with GNOVA, SBayesAPP provides finer resolution for identifying biologically meaningful annotations. Across the 15 T2D–trait analyses, SBayesAPP revealed clear tissue-specific enrichment patterns and a sharper separation between enriched and depleted annotations. Importantly, it not only detects enrichment but also attributes its source to effect-size correlation versus co-polygenicity. For example, in the T2D–fasting glucose analysis, enterocytes showed predominantly effect-size-driven enrichment, whereas erythrocytes showed polygenicity-driven enrichment. Together with simulations demonstrating approximately unbiased parameter estimates, these real-data results highlight SBayesAPP’s ability to prioritize T2D-relevant tissues and cell types and to distinguish the mechanisms underlying coheritability enrichment.

As noted by Lappalainen et al.^83^, polygenicity, like SNP-based heritability, can be partitioned across functional annotations. Boyle et al.^84^ showed that conserved regions are enriched for both heritability (13×) and polygenicity (14×), suggesting that their heritability enrichment largely arises from more causal variants rather than larger effects. This concept extends naturally to cross-trait architecture: within an annotation, coheritability enrichment arises from a corresponding enrichment in co-polygenicity. SBayesAPP, by jointly estimating co-polygenicity and SNP effect correlation, enables the dissection of the sources of coheritability enrichment. In the T2D–fasting glucose analysis, for instance, erythrocytes exhibited enrichment primarily driven by co-polygenicity rather than effect-size correlation. Given the essential role of erythrocytes in oxygen transport and their systemic impact on multiple organ systems, genetic variants that disrupt erythrocyte function are more likely to have widespread deleterious consequences. Such variants may therefore be subject to stronger purifying selection^85^, making large-effect pleiotropic variants rare and underrepresented in GWAS datasets^86^. This evolutionary constraint could result in enrichment being driven by many small-effect pleiotropic variants, reflected as high co-polygenicity with modest SNP effect correlation in erythrocytes.

Importantly, our results showed that coheritability enrichment is not always accompanied by polygenicity enrichment. While in some annotations a larger number of shared variants contributed to increased coheritability, others achieved enrichment primarily through stronger correlation of pleiotropic effect sizes. The ability of SBayesAPP to distinguish between polygenicity-driven enrichment and effect-size-driven enrichment enhances its interpretability and utility in characterizing the structure of shared genetic architecture.

Beyond estimating annotation-level SNP effect correlations and polygenic proportions, SBayesAPP computes SNP-level posterior probabilities, for each SNP and for each annotation it belongs to, of membership in four mixture polygenic components: null, pleiotropic, trait 1-specific, and trait 2-specific. These probabilities quantify, within a given annotation, the evidence that a SNP is shared across traits or specific to one trait. This information enables prioritization of candidate pleiotropic variants – something univariate analyses cannot resolve: running two single-trait models can flag variants for trait 1 and trait 2, but cannot distinguish a truly pleiotropic SNP from two distinct trait-specific causal SNPs in LD. Although LD can still blur assignments in the bivariate setting, modeling the cross-trait covariance helps reduce this ambiguity and improves discrimination. The reliability of SNP-level assignments depends on statistical power of data and on how distinct the annotation-specific mixture distributions are from each other. Greater separation of distributions across annotations yields more confident SNP assignments. A comprehensive evaluation of SBayesAPP’s SNP-level discovery performance is beyond the scope of this study and is left for future work.

SBayesAPP provides a powerful framework for dissecting shared genetic architecture, but performance can be limited by high levels of annotation overlap, which induces multicollinearity. Although SBayesAPP leverages annotation-specific mixture distributions to help resolve overlap, effectiveness depends on how distinct the annotation-stratified mixture distributions (including effect-size distributions and mixture components) are across annotations (e.g., based on biological separability). When a SNP belongs to multiple annotations that exhibit similar effect-size patterns and polygenic composition, the model may struggle to attribute its effect to the correct group. Thus, high-resolution, minimally overlapping annotations are essential for model’s identifiability and interpretability. Several additional limitations remain.

First, our analyses are restricted to individuals of European ancestry; future work should evaluate performance across diverse ancestral populations. Second, we focus on ∼1 million HapMap3 SNPs. As reported by Zheng et al.^4^, increasing genome coverage can improve polygenic prediction in univariate settings; extending SBayesAPP to higher-density arrays or whole-genome sequence data may reveal additional insights into shared genetic architecture.

Third, the cell-type annotations used in T2D analyses are derived from post-mortem tissues. Incorporating functional genomic data from live tissues or disease-relevant contexts could improve the biological relevance of annotation-based stratification. In addition, the anatomical origin of the single-cell references can bias results. In our schizophrenia and educational attainment analysis, brain cell-type annotations were derived primarily from midbrain samples, a region where inhibitory neurons are relatively more prominent in local circuitry than excitatory neurons. Thus, the depletion of heritability in excitatory neurons may reflect regional sampling bias rather than a true lack of involvement. Future work should integrate multi-region and stage-specific information to reduce such biases. Fourth, in our real-data comparisons of SBayesAPP and GNOVA, the uncertainty measures are not directly comparable. SBayesAPP reports posterior standard deviations from MCMC samples, whereas GNOVA reports frequentist standard errors obtained via the delta method. Although posterior standard deviations can approximate frequentist standard errors under large-sample regularity, cross-method comparisons of “significance” still need to be interpreted with caution.

In conclusion, SBayesAPP enables a biologically informed dissection of the shared genetic architecture underlying complex traits by jointly estimating annotation-stratified coheritability enrichment, SNP effect correlation, and polygenic proportions. This detailed level of insight into the sources of genetic correlation is not achievable with existing methods. Our results demonstrate that SBayesAPP produces accurate estimates of these annotation-specific parameters, offering meaningful interpretations of the biological mechanisms that contribute to trait etiology.

## Acknowledgements

This research was supported by the Australian National Health and Medical Research Council (1177268), and the Australian Research Council (DP220101947). J.Z. acknowledges support from Australian Research Council grant FL180100072 (awarded to P.M. Visscher). This study made use of data from the UK Biobank (project ID: 12505). H.C. acknowledges support from USDA–NIFA awards 2023-67015-39564 and 2023-70412-41054. J.Q. acknowledges support from USDA–NIFA award 2024-67034-42364.

## Author Contributions

H.C. and J.Z. conceived and supervised the study. J.Z., H.C., J.Q. and T.Z. developed the methods and algorithms. J.Z., H.C, J.Q. and T.Z. designed the experiment. J.Q. and T.Z. conducted all analyses with the assistance or guidance from T.L., A.L., S.L., S.C., L.Y., P.M.V., N.R.W., J.Z and H.C. J.Q., T.Z., J.Z., and H.C wrote the manuscript with the participation of all authors.

## Methods

### Summary-data-based low-rank model

In this section, we outline a summary-data-based single-trait model derived from the general form of a multiple regression model using individual-level data, as proposed by Zheng et al.^1^ Consider the following individual-data-based model:

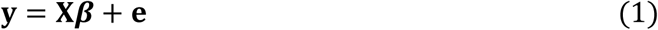

where **y** is the vector of trait phenotypes adjusted for covariates; **X** is the genotype matrix for m SNPs, standardized with mean zero and unit variance; 𝜷 is the vector of true SNP effects, with residual variance being 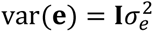. Let N be the sample size of a GWAS analysis, we multiply both sides of equation (1) by 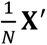 to obtain:

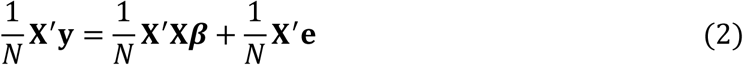

with the left-hand side being the conventional GWAS marginal effect estimates 𝐛. If we consider 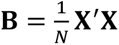 as the LD correlation matrix, we can rewrite equation (2) as a standard summary-data-based model:

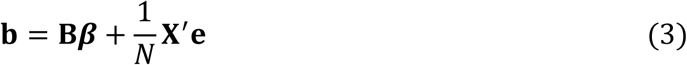

with residual variance 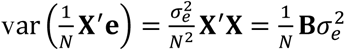

Due to the hierarchical LD structure of the human genome, approximately independent LD blocks have been identified across human populations^2^. The sub-matrices of the whole-genome LD matrix corresponding to each LD block can be considered approximately independent. Hence, the genome-wide LD matrix can be represented as a block-diagonal matrix with blocks defined by LD blocks. This structure allows the summary-data-based model to be divided into smaller, independent models based on LD blocks. SNP effects within each LD block can then be estimated in parallel, enabling more efficient computation. To succinctly illustrate the independence between LD blocks, consider a simple example. Assuming the whole genome is composed of three LD blocks, equation (3) can be further expressed as:

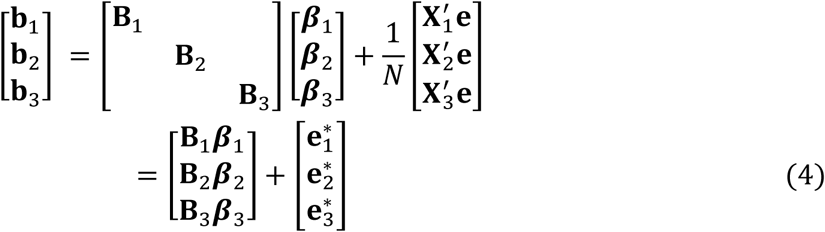

with

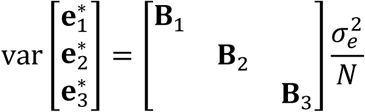

Due to the independence across blocks, the estimation of 𝜷_i_ within an LD block i can be performed independently of the estimations within other blocks.

Although SNP effects are estimated independently within LD blocks in summary-data-based model, global parameters, such as the SNP effect (co)variance matrix and the polygenic mixture proportions, should be estimated genome-wide. Performing this genome-wide inference is computationally intensive, particularly in a Bayesian framework. To address this challenge, Zheng et al.^1^ proposed a low-rank model that enhances effect sampling by leveraging the eigen-decomposition of the block LD matrix. In each LD block, the LD matrix can be decomposed as:

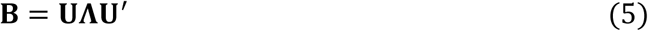

where **U** is the matrix of eigenvectors and **Λ** is the diagonal matrix of eigenvalues. Substituting **B** in equation (3) yields:

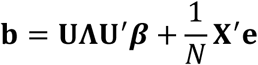

Multiplying both sides by

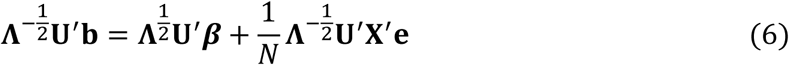

Equation (6) can be simplified as 𝐰 = 𝐐𝜷 + 𝝐, where

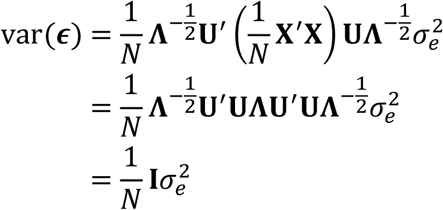

The LD matrix estimated from a reference panel often exhibits rank deficiency due to LD between SNPs and the limitations of sample size. In such cases, several eigenvalues become zero. Additionally, small eigenvalues are particularly susceptible to sampling variations in LD between the GWAS and the reference samples. Therefore, we divided 𝚲 as:

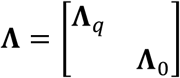

where 𝚲_𝑞_ contains the top 𝑞 eigenvalues arranged in descending order, explaining at least a predefined proportion of variance (e.g., 99.5%) in the LD matrix. The remaining eigenvalues, including zeros, are stored in 𝚲_0_. Equation (6) can be further expressed as:

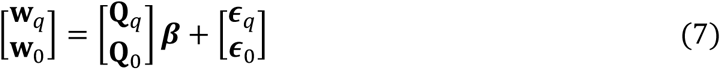

To remove noise in the LD matrix, we discard the equations associated with 𝐰_0_, resulting in a low-rank model:

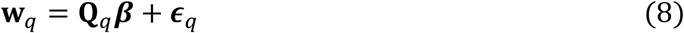

with

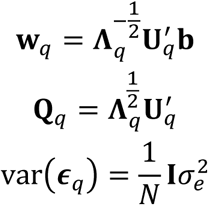

#### Summary-data-based low-rank two-trait model

The derivation process described in the previous section leads us to the development of a summary-data-based low-rank two-trait model. Assuming that GWAS for two traits are conducted within the same population, the summary-data-based model for these two traits within a single LD block can be expressed as follows:

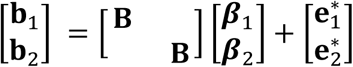

where 𝐛_𝑡_ denotes the marginal SNP effects obtained from single-trait GWAS analysis for trait 𝑡, 𝐁 denotes the LD matrix for the corresponding LD block, and 𝜷_𝑡_ denotes the true SNP effects for trait 𝑡. The variance of the residuals is given by 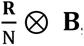, where 𝐑 is the residual variance-covariance matrix across traits. After the eigen-decomposition of 𝐁 and its subsequent substitution into equation 8, we obtain the low-rank two-trait model:

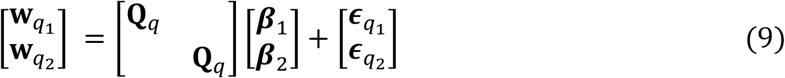

The residual variance matrix for the summary-data-based low-rank two-trait model (equation 9) is expressed as 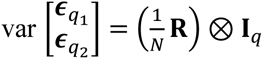, where 𝑞 denotes the number of top eigenvalues that explain at least a predefined proportion of variance in the LD matrix.

Notably, the structure of the summary-data-based low-rank two-trait model closely parallels that of the individual-level-based model^3^. In this framework, the phenotypic vector for trait 𝑡 in the individual-data-based model corresponds to 𝐰_𝑞𝑡_ in the low-rank model, while the genotypic covariate (design) matrix corresponds to 𝐐_𝑞_. Moreover, the residual variance remains independent within each trait for both models, with the variance-covariance matrix across traits in the low-rank model scaled by a factor of 𝑁 relative to the individual-data-based model. As a result, Gibbs sampling methods for the summary-data-based low-rank two-trait model closely resemble those used in the individual-data-based two-trait model^3^.

#### Low-rank two-trait model incorporating functional annotation and mixture prior (SBayesAPP)

Here, we further integrate functional annotations into the low-rank two-trait model employing mixture priors for each annotation, hereafter referred to as SBayesAPP. In this model, we assume that the total effect of each SNP is partitioned into distinct components corresponding to different annotation groups. Specifically, given a functional annotation matrix 𝐀 of dimension 𝑚 × 𝑐 (where 𝑚 is the number of SNPs and 𝑐 is the number of annotation groups), the SNP effects at locus j are modeled as:

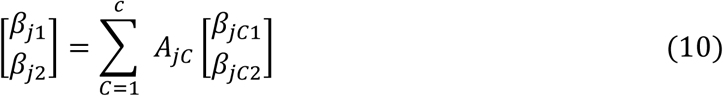

where 𝛽_𝑗𝑡_ denotes the total effect of SNP 𝑗 on trait 𝑡, 𝛽_𝑗𝐶𝑡_ denotes the partial effect of SNP 𝑗 on trait 𝑡 attributable to annotation 𝐶, and 𝐴_𝑗𝐶_ denotes the annotation covariate for SNP 𝑗 associated with annotation 𝐶. For categorical annotation groups, 𝐴_𝑗𝐶_ is binary (taking values of 0 or 1), whereas for continuous annotation groups, 𝐴_𝑗𝐶_ corresponds to the observed values. For each pair of SNP effects at locus 𝑗 corresponding to annotation 𝐶, a two-trait BayesC mixture prior is assigned. The annotation-specific SNP effect variance-covariance matrix and annotation-specific polygenic proportions are considered unknown and are estimated during model fitting. The distribution of annotation-specific SNP effects for locus j in annotation 𝐶 is given by:

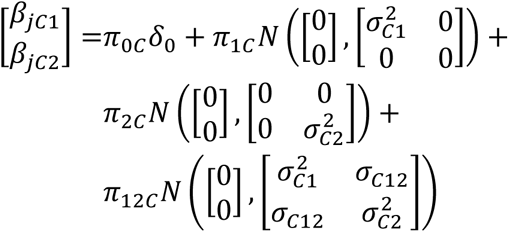

where 𝛿_0_denotes a point mass at 0, corresponding to null-effect SNPs. The SNP effect variance for trait 𝑡 in annotation 𝐶 is represented by 𝜎_Ct_^2^, and 𝜎_𝐶12_ represents the covariance of SNP effects within annotation 𝐶. The annotation-specific genetic architecture is characterized by: 𝜋_0𝐶_, the proportion of null-effect SNPs; 𝜋_1𝐶_, the proportion of SNPs influencing only trait 1; 𝜋_2𝐶_, the proportion of SNPs influencing only trait 2, and 𝜋_12𝐶_, the proportion of pleiotropic SNPs affecting both traits.

Through the proposed framework of SBayesAPP, each trait is modeled using 𝑐 multiple regressions, corresponding to 𝑐 different annotation groups, with the regression coefficients weighted by their respective annotation covariates. To illustrate this, consider an example with two annotation groups and three SNPs. The annotation matrix 𝐀 is defined as:

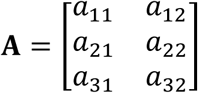

where 𝑎_𝑗𝐶_ represents the annotation covariate of SNP 𝑗 in annotation 𝐶. The total SNP effects (β) of the three SNPs for one trait can be expressed as:

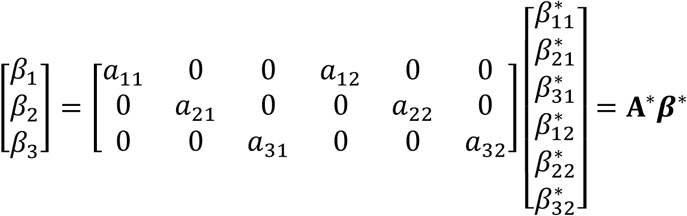

where 𝐀^∗^is the transformed annotation matrix, and 𝜷^∗^is a vector of annotation-specific SNP effects, arranged by annotation groups. Specifically, 𝛽^∗^_*jc*_ represents the effect of SNP 𝑗 attributed to annotation 𝐶. With the integration of functional annotation, the low-rank model described in equation (8) can be extended as:

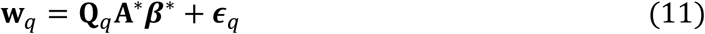

This model can be decomposed into 𝑐 components, corresponding to the 𝑐 annotation groups:

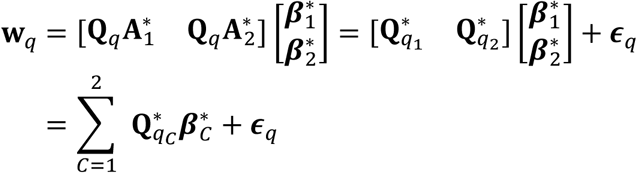

where 𝐀^∗^_*C*_ is a diagonal matrix with the diagonal elements being the covariates for annotation 𝐶. Thus, the two-trait model can be expressed as:

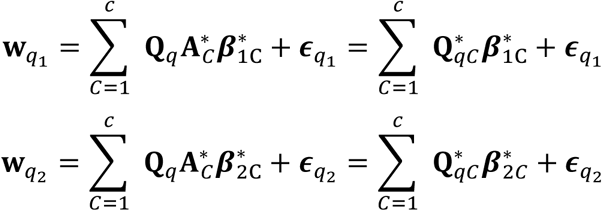

#### Full conditional distribution of annotation-specific effects

The full conditional distribution for the annotation-specific effect of SNP 𝑗 in annotation 𝐶 is given by

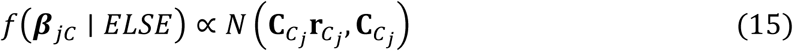

where 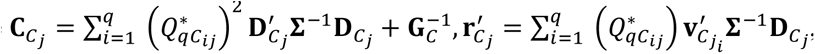, and 𝐯_𝐶𝑗1_ = 𝐰*_i_* – 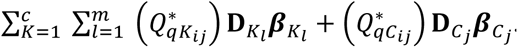. Full derivations of the Gibbs sampler are provided in the Supplementary File. In the low-rank two-trait model, each block-specific residual (co)variance matrix is assigned an inverse-Wishart prior (see Supplementary File). Each annotation-specific SNP effect (co)variance matrix (𝐆_C_) likewise receives an inverse-Wishart prior. The annotation-stratified polygenic proportions, Π_C_ =[𝜋_0𝐶_, 𝜋_1𝐶_, 𝜋_2𝐶_, 𝜋_12𝐶_], are given a uniform Dirichlet prior.

#### Coheritability Enrichment as a Measure of Annotation Importance

In this study, we use coheritability (co-h^2^) enrichment to assess whether the genetic covariance between two traits is disproportionately explained by variants within a specific annotation group, relative to genome-wide expectations. This metric compares the per-SNP genetic covariance within an annotation to the per-SNP genome-wide genetic covariance, effectively normalizing for annotation size and enabling fair comparisons across functional categories. In contrast, annotation-specific genetic correlation quantifies the genetic correlation between traits within a specific annotation. However, it is influenced not only by genetic covariance but also by the heritability (h^2^) of each trait, making it less informative for determining whether an annotation is overrepresented in shared genetic architecture. Notably, a high genetic correlation within an annotation does not necessarily imply a large contribution to the overall genetic covariance.

Therefore, compared to genetic correlation, coheritability enrichment offers a more robust and biologically interpretable measure for prioritizing functionally relevant annotations. It ensures that the estimated genetic contribution reflects meaningful patterns of shared genetic architecture, without being confounded by annotation size or trait-specific heritability.

#### Simulation

We simulated datasets using real genotypes from 300,000 individuals of European ancestry, obtained from the UK Biobank^6^ (UKB). Quality control followed procedures described in Zheng et al.¹. Simulations were performed using 95,782 SNPs from chromosome 1, based on the 1M HapMap3 SNP set. We modeled two genetically correlated traits, with heritability values ℎ_1_^2^ = 0.5 (Trait 1) and ℎ^2^ = 0.2 (Trait 2). SNPs were divided into two non-overlapping annotation groups based on retinal bipolar cell type annotations: 9% of SNPs were assigned to annotation 1 (𝐶1), and the remaining 91% to annotation 2 (𝐶2). Within each annotation, 1% of SNPs were randomly selected as causal variants.

We simulated four levels of co-polygenicity among causal variants (𝑝), specifically 𝑝 = 1, 0.8, 0.5, and 0.1, indicating the proportion of causal variants that affected both traits. When 𝑝 = 1, all causal variants were pleiotropic. For 𝑝 < 1, a fraction 𝑝 of causal variants in each annotation group were pleiotropic, while the remaining 1 − 𝑝 were equally split to affect only Trait 1 or

Trait 2. To introduce coheritability enrichment, we ensured that each annotation group explained 50% of the total heritability for both traits, regardless of annotation size. SNP effect correlations were defined as 0.8 for 𝐶1 and 0.2 for 𝐶2, introducing differential coheritability across annotation groups. Since the SNP effect covariance matrix depends on co-polygenicity, annotation size, annotation-specific heritability, and SNP effect correlation, we first computed the annotation-specific genetic correlation as:

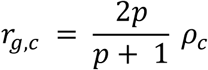

where 𝜌_𝑐_ is the predefined SNP effect correlation for annotation 𝑐. The SNP effect covariance matrix for each annotation was constructed as:

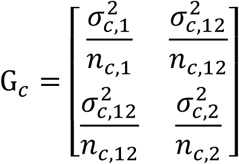

where 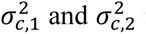 represent the ℎ^2^ contributions of annotation 𝑐 to Trait 1 and Trait 2, respectively, both set to half of the corresponding total heritability 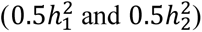. The term 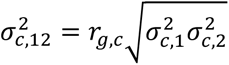 captures the genetic covariance. Here, 𝑛*_c,_*_1_ and 𝑛*_c,_*_2_ denote the total number of causal variants affecting Trait 1 and Trait 2 within annotation 𝑐, respectively, while 𝑛_𝑐,12_ denotes the number of pleiotropic variants within annotation 𝑐. Based on the simulated SNP effects, polygenic risk scores (PRS) for each trait were computed as:

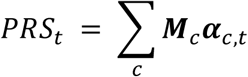

where 𝑴_𝑐_ is the genotype matrix for causal variants in annotation 𝑐, and 𝜶_𝑐,𝑡_ represents their effect sizes for trait 𝑡. Residuals were sampled from a bivariate normal distribution with residual correlation 𝜌_𝑟𝑒𝑠_ = 0.1, and the residual covariance matrix was formulated as:

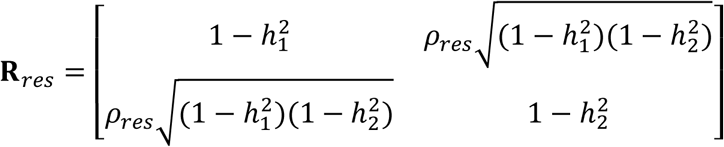

Standardized phenotypes for each trait were then generated by summing PRS and residuals. For each level of co-polygenicity among causal variants (𝑝 *=* 1, 0.8, 0.5, 0.1), we generated 20 replicate datasets.

Empirically, the average annotation-specific genetic correlations across the 20 replicates, derived from the predefined SNP effect correlations (0.8 for 𝐶1 and 0.2 for 𝐶2) were 0.79, 0.70, 0.53, and 0.16 for 𝐶1 and 0.19, 0.17, 0.14, and 0.04 for 𝐶2, under 𝑝 *=* 1, 0.8, 0.5, and 0.1, respectively. These results illustrate that genetic correlation alone cannot fully reflect the strength of pleiotropy, as it is also influenced by annotation size and the proportion of trait-specific (non-pleiotropic) variants. GWAS summary statistics were generated using PLINK^9^.

The performance of the SBayesAPP in simulation studies was evaluated by comparing its estimates of key genetic parameters, including annotation-specific genetic correlation, co-ℎ^2^enrichment, ℎ^2^enrichment, SNP effect correlation, and polygenicity among causal variants, to the corresponding true simulated values. To obtain starting values for the SBayesAPP model, single-trait BayesC mixture models were first applied to estimate trait-specific heritability and polygenicity. These estimates were then used to initialize the annotation-specific polygenic proportions and the SNP effect (co)variance matrix. SBayesAPP was run for 1,000 MCMC iterations per replicate, with the first 400 iterations treated as burn-in. During the burn-in period, the scale parameter of the prior distribution for the annotation-specific SNP effect covariance matrix was estimated. In addition, block-specific residual (co)variance matrices were estimated for each LD block and subsequently used in the Gibbs sampling of SNP effects within the corresponding block. SNP effect correlations were derived from the (co) variance matrices of pleiotropic SNPs.

Increasing the MCMC chain length beyond 1,000 iterations did not improve estimation accuracy in the simulation analysis. For comparison, results from LDSC, HDL, SLDSC, and GNOVA were also evaluated under each simulation setting, based on the genetic parameters that each model is capable of estimating.

#### Real data analysis

All real data analyses used ∼1 million SNPs from the HapMap3 reference panel. For the cell type analyses, we used genomic cell-type annotations that were generated using Expression Weighted Cell Type Enrichment (EWCE) method^7^. Three single-cell resources were used: (i) single-cell transcriptomic data (protein coding genes only) from Tabula Sapiens Consortium^4^ (TS) for the T2D-related analyses, (ii) brain substantia nigra pars compacta (SNpc)^5^ for the analysis between schizophrenia and educational attainment, and (iii) Human Lung Cell Atlas — core healthy lung parenchyma^6^ (LP) for the analysis between smoking and lung cancer. These resources provided 162 TS cell-type annotations, 8 SNpc major cell-type annotations, and 42 LP cell-type annotations, respectively. For the T2D-related analyses, 155 of the 162 TS annotations contained at least one HapMap3 SNP; accordingly, we analyzed those 155 annotations plus a background group comprising SNPs not assigned to any of the 155 annotations. A heatmap of Jaccard indices quantifying SNP overlap among the 155 cell type annotations is shown in **Supplementary Figure 5**.

The proof-of-principle analysis was conducted using a single MCMC chain with 3,000 iterations. For the rest of cell type analyses, SBayesAPP was run for 3,000 iterations across 10 independent chains to ensure stable and robust estimation. Extending the chain length beyond 3,000 iterations did not meaningfully alter the annotation-specific enrichment estimates within individual chains (see **Supplementary Figure 6**). The first 1000 iterations were treated as burn-in. During the initial 2,000 iterations, the scale parameter of the prior distribution for the SNP effect covariance matrix was estimated and then fixed for the remaining iterations. Residual variances were sampled in a block-wise manner. We assessed the convergence of coheritability enrichment results by examining the correlations of enrichment estimates across grouped chains (see **Supplementary Figure 6**). Posterior means and standard deviations were computed from the MCMC samples of the ten-chain means. For comparison, we also ran GNOVA using default settings and computed coheritability enrichment based on genetic covariance estimates corrected for sample overlap.

## Data Availability

The UK Biobank (UKB) dataset used in simulation study can be accessed through formal application procedures detailed at the UKB website (http://www.ukbiobank.ac.uk). Summary statistics for all the traits utilized in real data analysis are available from their original publications. Additionally, the eigen-decomposition data for linkage disequilibrium (LD) matrices are accessible via the Genome-wide Complex Trait Bayesian (GCTB) software website at https://gctbhub.cloud.edu.au/software/gctb/#Download.

## Code Availability

SBayesAPP software is publicly available and implemented in the Julia programming language, with source code and documentation hosted at https://github.com/reworkhow/SBayesAPP/. All analysis scripts used for this manuscript are openly available at: https://github.com/reworkhow/SBayesAPP/.

