## Supplementary File for "Quantifying annotation-stratified pleiotropy and co-polygenicity between complex traits"

### Supplementary File for SBayesAPP

#### 1 Derivation of the Gibbs sampler for SNP-specific effects

$$\mathbf{w}_1 = \sum_{C=1}^{c} \mathbf{Q}_C \boldsymbol{\beta}_{1C} + \boldsymbol{\epsilon}_1$$
$$\mathbf{w}_2 = \sum_{C=1}^{c} \mathbf{Q}_C \boldsymbol{\beta}_{2C} + \boldsymbol{\epsilon}_2$$

For element $i$, with multi-trait BayesC mixture prior implemented, its likelihood function is shown as

$$\mathbf{w}_i = \sum_{C=1}^{c} \sum_{j=1}^{m} Q_{C_{ij}} \mathbf{D}_{C_j} \boldsymbol{\beta}_{C_j} + \boldsymbol{\epsilon}_i$$

where $\mathbf{w}_i$ is a $2 \times 1$ vector corresponding to the $i$th element of the trait-pair vectors $\mathbf{w}_1$ and $\mathbf{w}_2$; $Q_{C_{ij}}$ denotes the $(i,\ j)$th element of the annotation-specific design matrix $\mathbf{Q}_C$, $\mathbf{D}_{C_j}$ is a $2 \times 2$ diagonal matrix indicating whether the SNP at locus $j$ has no effect, an effect on one trait, or effects on both traits under annotation $C$, $\boldsymbol{\beta}_{C_j}$ is a $2 \times 1$ vector of SNP effect sizes at locus $j$ for traits associated with annotation $C$; and the residual variance is given by $var(\boldsymbol{\epsilon}_i) = \boldsymbol{\Sigma} = \frac{1}{N}\boldsymbol{R}$, where N is the sample size and $\mathbf{R}$ is the covariance matrix across traits. The full conditional distribution of $\boldsymbol{\beta}_{C_j}$ is derived as

$$
\begin{aligned}
f(\boldsymbol{\beta}_{C_j}|ELSE) &\propto f(\mathbf{w}|ELSE)f(\boldsymbol{\beta}_{C_j}|\mathbf{G}_C) \\
&\propto \exp\left[-\sum_{i=1}^{q}\frac{(\mathbf{v}_{Cj_i}-Q_{C_{ij}}\mathbf{D}_{C_j}\boldsymbol{\beta}_{C_j})'\boldsymbol{\Sigma}^{-1}(\mathbf{v}_{Cj_i}-Q_{C_{ij}}\mathbf{D}_{C_j}\boldsymbol{\beta}_{C_j})}{2}\right]\times\exp\left(-\frac{1}{2}\boldsymbol{\beta}'_{C_j}\mathbf{G}_C^{-1}\boldsymbol{\beta}_{C_j}\right) \\
&\propto \exp\left[-\frac{1}{2}\sum_{i=1}^{q}(\mathbf{v}'_{Cj_i}\boldsymbol{\Sigma}^{-1}\mathbf{v}_{Cj_i}-2Q_{C_{ij}}\mathbf{v}'_{Cj_i}\boldsymbol{\Sigma}^{-1}\mathbf{D}_{C_j}\boldsymbol{\beta}_{C_j}+Q_{C_{ij}}^2\boldsymbol{\beta}'_{C_j}\mathbf{D}'_{C_j}\boldsymbol{\Sigma}^{-1}\mathbf{D}_{C_j}\boldsymbol{\beta}_{C_j})\right] \\
&\times\exp\left(-\frac{1}{2}\boldsymbol{\beta}'_{C_j}\mathbf{G}_C^{-1}\boldsymbol{\beta}_{C_j}\right) \\
&\propto \exp\{-\frac{1}{2}[\boldsymbol{\beta}'_{C_j}(\sum_{i=1}^{q}Q_{C_{ij}}^2\mathbf{D}'_{C_j}\boldsymbol{\Sigma}^{-1}\mathbf{D}_{C_j}+\mathbf{G}_C^{-1})\boldsymbol{\beta}_{C_j}-2\sum_{i=1}^{q}Q_{C_{ij}}\mathbf{v}'_{Cj_i}\boldsymbol{\Sigma}^{-1}\mathbf{D}_{C_j}\boldsymbol{\beta}_{C_j}]\} \\
&\propto \exp\{-\frac{1}{2}[\boldsymbol{\beta}'_{C_j}\mathbf{C}_{C_j}\boldsymbol{\beta}_{C_j}-2\mathbf{r}'_{C_j}\boldsymbol{\beta}_{C_j}]\} \\
&\propto \exp\{-\frac{1}{2}(\boldsymbol{\beta}'_{C_j}-\mathbf{r}'_{C_j}\mathbf{C}_{C_j}^{-1})\mathbf{C}_{C_j}(\boldsymbol{\beta}_{C_j}-\mathbf{C}_{C_j}^{-1}\mathbf{r}_{C_j})\} \\
&\propto N(\mathbf{C}_{C_j}^{-1}\mathbf{r}_{C_j},\mathbf{C}_{C_j}^{-1})
\end{aligned}
$$

where $\mathbf{C}_{C_j} = \sum_{i=1}^{q} Q_{C_{ij}}^2\mathbf{D}'_{C_j}\boldsymbol{\Sigma}^{-1}\mathbf{D}_{C_j}+\mathbf{G}_C^{-1}$, $\mathbf{r}'_{C_j} = \sum_{i=1}^{q} Q_{C_{ij}}\mathbf{v}'_{Cj_i}\boldsymbol{\Sigma}^{-1}\mathbf{D}_{C_j}$, and $\mathbf{v}_{Cj_i} = \mathbf{w}_i - \sum_{K=1}^{c}\sum_{l=1}^{m} Q_{K_{il}}\mathbf{D}_{K_l}\boldsymbol{\beta}_{K_l} + Q_{C_{ij}}\mathbf{D}_{C_j}\boldsymbol{\beta}_{C_j}$.

### 2 Thresholding implementation and block-wise Gibbs sampler for residual variance in real data analysis

#### 2.1 Residual (co)variance matrix for summary-data-based low-rank two-trait model when GWAS sample sizes are not consistent

The summary-statistics-based model for two traits within a single LD block can be expressed as:

$$\mathbf{b}_1 = \mathbf{B}\boldsymbol{\beta}_1 + \frac{1}{N_1}\mathbf{X}'\mathbf{e}_1 = \mathbf{B}\boldsymbol{\beta}_1 + \mathbf{e}_1^* \tag{1}$$

$$\mathbf{b}_2 = \mathbf{B}\boldsymbol{\beta}_2 + \frac{1}{N_2}\mathbf{X}'\mathbf{e}_2 = \mathbf{B}\boldsymbol{\beta}_2 + \mathbf{e}_2^* \tag{2}$$

Equation 1 assumes that the sample size of the LD reference population is equivalent to the GWAS sample size of trait 1 (i.e., $N_1$) with $var(\mathbf{e}_1^*) = \frac{1}{N_1}\mathbf{B}\sigma_{e_1}^2$, and equation 2 assumes that the sample size of the LD reference population is equivalent to the GWAS sample size of trait 2 (i.e., $N_2$) with $var(\mathbf{e}_2^*) = \frac{1}{N_2}\mathbf{B}\sigma_{e_2}^2$. After eigen-decomposition, the low-rank two-trait model derived from equation 1 and 2 can be expressed as

$$\begin{aligned}\mathbf{w}_1 &= \mathbf{Q}\boldsymbol{\beta}_1 + \frac{1}{N_1}\boldsymbol{\Lambda}^{-\frac{1}{2}}\mathbf{U}'\mathbf{X}'\mathbf{e}_1 = \mathbf{Q}\boldsymbol{\beta}_1 + \boldsymbol{\epsilon}_1 \\ \mathbf{w}_2 &= \mathbf{Q}\boldsymbol{\beta}_2 + \frac{1}{N_2}\boldsymbol{\Lambda}^{-\frac{1}{2}}\mathbf{U}'\mathbf{X}'\mathbf{e}_2 = \mathbf{Q}\boldsymbol{\beta}_1 + \boldsymbol{\epsilon}_2\end{aligned}$$

where $var(\boldsymbol{\epsilon}_1) = \frac{1}{N_1}\mathbf{I}\sigma_{e_1}^2$ and $var(\boldsymbol{\epsilon}_2) = \frac{1}{N_2}\mathbf{I}\sigma_{e_2}^2$. Then,

$$\begin{aligned}cov(\boldsymbol{\epsilon}_1, \boldsymbol{\epsilon}_2) &= cov((\frac{1}{N_1}\boldsymbol{\Lambda}^{-\frac{1}{2}}\mathbf{U}'\mathbf{X}'\mathbf{e}_1), (\frac{1}{N_2}\boldsymbol{\Lambda}^{-\frac{1}{2}}\mathbf{U}'\mathbf{X}'\mathbf{e}_2)) \\ &= \frac{1}{N_1N_2}\boldsymbol{\Lambda}^{-\frac{1}{2}}\mathbf{U}'\mathbf{X}'cov(\mathbf{e}_1, \mathbf{e}_2)\mathbf{X}\mathbf{U}\boldsymbol{\Lambda}^{-\frac{1}{2}}\end{aligned}$$

Since covariance $\sigma_{12}$ only exists on the $N_B$ overlapping individuals, and we assume the LD matrix is stable, i.e., $\mathbf{X}_B'\mathbf{X}_B \approx N_B\mathbf{B}$ with $\mathbf{B}$ being the LD matrix, we have

$$\begin{aligned}cov(\boldsymbol{\epsilon}_1, \boldsymbol{\epsilon}_2) &= \frac{\sigma_{12}}{N_1N_2}\boldsymbol{\Lambda}^{-\frac{1}{2}}\mathbf{U}'\mathbf{X}_B'\mathbf{X}_B\mathbf{U}\boldsymbol{\Lambda}^{-\frac{1}{2}} \\ &= (\frac{N_B}{N_1N_2}\sigma_{12})\boldsymbol{\Lambda}^{-\frac{1}{2}}\mathbf{U}'\mathbf{U}\boldsymbol{\Lambda}\mathbf{U}'\mathbf{U}\boldsymbol{\Lambda}^{-\frac{1}{2}} \\ &= \frac{N_B}{N_1N_2}\sigma_{12}\mathbf{I}\end{aligned}$$

Therefore, the residual (co)variance matrix for the low-rank two-trait model is

$$\mathbf{\Sigma} = \begin{bmatrix} \frac{1}{N_1}\mathbf{I}\sigma_1^2 & \frac{N_B}{N_1 N_2}\mathbf{I}\sigma_{12} \\ \frac{N_B}{N_1 N_2}\mathbf{I}\sigma_{12} & \frac{1}{N_2}\mathbf{I}\sigma_2^2 \end{bmatrix} \tag{3}$$

In our MCMC framework, the $2 \times 2$ residual covariance matrix $\mathbf{\Sigma}$ is given an inverse-Wishart prior and updated block-wise across the genome. Specifically, for each LD block $b$, we sample a block-specific residual covariance $\mathbf{\Sigma}_b$ at every iteration. The prior on $\mathbf{\Sigma}_b$ is specified as:

$$\mathbf{\Sigma}_{prior} \sim \mathcal{IW}(\nu_0, \mathbf{S}_0)$$

where $\nu_0 = 4 + n_{trait}$ and $\mathbf{S}_0 = \text{diag}(\frac{1}{N_{1,ave}}, \frac{1}{N_{2,ave}}) \cdot (\nu_0 - n_{trait} - 1)$, corresponding to a weakly informative identity prior for the bivariate residual variances. Here, $N_{1,ave}$ and $N_{2,ave}$ are the average sample sizes for trait 1 and trait 2, respectively. At each iteration, the full conditional posterior for the residual covariance matrix in block $b$ is:

$$\mathbf{\Sigma}_b \sim \mathcal{IW}(\nu_0 + n_b, \mathbf{S}_0 + \mathbf{SSE}_b)$$

where $n_b$ denotes the block-specific number of response variables (i.e., number of eigenvectors), and $\mathbf{SSE}_b$ is the cross-product matrix of the block-specific residuals $\epsilon_{1,b}$ and $\epsilon_{2,b}$.

To enforce consistency between genetic and residual variance, we apply a thresholding heuristic to the diagonal elements $\Sigma_{b,ii}$ of the block-specific residual covariance. For each trait $i$, define

$$\text{thres}_{b,i} = \frac{\sum_C \alpha_{b,i}^2}{(\mathbf{Q}\boldsymbol{\alpha}_i)^T(\mathbf{Q}\boldsymbol{\alpha}_i)}$$

where $\sum_C \alpha_{b,i}^2$ is the sum of squared SNP effects across the $C$ annotations for trait $i$, and $(\mathbf{Q}\boldsymbol{\alpha}_i)^T(\mathbf{Q}\boldsymbol{\alpha}_i)$ is the genetic variance of the polygenic score for trait $i$ in block $b$. If $\text{thres}_{b,i} > 1.1$ ( i.e., the SNP-effect estimates explain more variance than the polygenic scores), the sampled residual variance $\Sigma_{b,ii}$ is retained; otherwise, $\Sigma_{b,ii}$ is fixed to $\frac{1}{N_{i,ave,b}}$

(assuming standardized phenotypes), where $N_{i,ave,b}$ is the block-specific average sample size. This thresholding guards against inflated or underestimated residual variances in poorly identified blocks. The off-diagonal elements of $\mathbf{\Sigma}_b$ are then reconstructed from the sampled correlation:

$$\Sigma_{b,12} = \Sigma_{b,21} = \rho_\Sigma \cdot \sqrt{\Sigma_{b,11} \cdot \Sigma_{b,22}}$$

where $\rho_\Sigma$ is the residual correlation computed from the sampled $\mathbf{\Sigma}_b$ before thresholding.

### 3 Empirical estimation of prior scale for SNP effect covariance matrix

To estimate the scale parameter of the inverse-Wishart prior used for annotation-stratified SNP effect covariance matrices, we performed an empirical update of the prior during the first 2000 iterations. Specifically, we estimated a correlated SNP effect covariance matrix $\mathbf{G}_{\text{prior}} \in \mathbb{R}^{2\times 2}$ for each annotation group by aggregating information from the MCMC samples of SNP effects. At each iteration, we computed a sum-of-squares matrix (denoted **SSQ**) using the dot product of sampled SNP effects across all SNPs in a given annotation for each pair of traits. This matrix includes both diagonal elements (trait-specific variances) and off-diagonal elements (cross-trait covariances). To ensure stability, we accumulated a moving average of these matrices over iterations, denoted **meanSSQ**. The resulting averaged matrix is then divided by the number of SNPs in the annotation group to yield an empirical (co)variance matrix estimate:

$$\mathbf{G}_{\text{prior},c} = \frac{\mathbf{meanSSQ}_c}{n_c}$$

where $n_c$ is the number of SNPs in annotation $c$. Finally, this matrix is scaled by $(\text{df}_G - n_{trait} - 1)$ to produce the prior scale matrix used in the inverse-Wishart distribution for that annotation at current iteration. This empirical approach allows the model to adaptively learn appropriate levels of trait-specific and cross-trait sharing in SNP prior distributions, stratified by functional annotations.

#### 4 Definition and Computation of Annotation Enrichment in SBayesAPP

**Definition of Enrichment** Enrichment quantifies whether a specific set of genetic variants (e.g., in a functional genomic region) explains more heritability (or co-heritability) than expected under a null hypothesis of uniform contribution per SNP. Formally, the enrichment of annotation $A$ can be defined as

$$\text{Enrichment}(A) = \frac{\text{per-SNP } h^2 \text{ (or co}h^2\text{) explained by SNPs in } A}{\text{per-SNP } h^2 \text{ (or co}h^2\text{) expected under the null}}$$

Under the null hypothesis, every SNP contributes equally to the total $h^2$ (or co-$h^2$), so the expected per-SNP contribution is $\frac{h^2_{\text{total}}}{n_{\text{SNP, total}}}$. In SBayesAPP, each SNP's effect is decomposed additively across all annotations it belongs to. Accordingly, if a SNP is in $c$ overlapping annotations, the null contribution is split evenly across them: the per-annotation null contribution for the SNP is $\frac{1}{c}\frac{h^2_{\text{total}}}{n_{\text{SNP, total}}}$.

**Example** Consider annotation $A_1$, which includes: $n_1 - n_{12}$ SNPs unique to $A_1$, and $n_{12}$ SNPs shared with $A_2$. under the null hypothesis, the expected heritability attributed to $A_1$ is:

$$h^2_{A_1,\text{null}} = \frac{h^2_{\text{total}}}{n_{\text{SNP, total}}}\left[(n_1 - n_{12}) + \tfrac{1}{2}n_{12}\right].$$

Therefore, the enrichment for $A_1$ is

$$\text{Enrichment}(A_1) = \frac{h^2_{A_1}}{h^2_{A_1,\text{null}}} = \frac{\frac{h^2_{A_1}}{(n_1 - n_{12} + \frac{1}{2}n_{12})}}{\frac{h^2_{\text{total}}}{n_{\text{SNP, total}}}}.$$

**General Formula** For a general annotation $A$, define the overlap-adjusted SNP count $n_A^{adj}$ as

$$n_A^{adj} = \sum_{i \in A} \frac{1}{c_i},$$

where $c_i$ is the number of annotations that SNP $i$ belongs to. This adjustment accounts for overlapping annotations by allocating each SNP's contribution equally across the annotation categories it belongs to. The expected heritability explained by annotation A under the null hypothesis is then

$$h^2_{A,\text{null}} = \frac{h^2_{\text{total}}}{n_{\text{SNP, total}}} n_A^{adj},$$

and the enrichment is computed as

$$\text{Enrichment}(A) = \frac{h^2_A}{h^2_{A,\text{null}}} = \frac{\frac{h^2_A}{n_A^{adj}}}{\frac{h^2_{\text{total}}}{n_{\text{SNP, total}}}}.$$

This formulation naturally extends to co-heritability (co$h^2$) by replacing $h^2_A$ and $h^2_{\text{total}}$ with the corresponding co-heritability estimates.
