## Supplementary Table and Figure for "Quantifying annotation-stratified pleiotropy and co-polygenicity between complex traits"

**Supplementary File**

**Supplementary Table 1.** Summary of all the analyzed traits.

| Trait name | Acronym | Sample size^1^ | Reference | $h^{2}$^2^ | $r_{g}$ with T2D^3^ | Number of COJO SNPs^4^ |
| --- | --- | --- | --- | --- | --- | --- |
| Atrial fibrillation | AF | 1,030,840 | Nielsen, Jonas B., et al. Nature genetics (2018) | 0.029 | -0.029 | 152 |
| Body mass index | BMI | 685,723 | Yengo, Loic, et al. 2018 | 0.217 | 0.445 | 1021 |
| Cholesterol | CHOL | 318,927 | Sinnott-Armstrong, Nasa, et al. Nature genetics (2021) | 0.245 | -0.018 | 414 |
| Depression | DEP | 500,199 | Howard, David M., et al. Nature neuroscience (2019) | 0.069 | 0.028 | 108 |
| Educational attainment | EA | 766,345 | Okbay, Aysu, et al.  Nature genetics (2022) | 0.117 | -0.144 | 470 |
| Fasting glucose | FG | 88,971 | Lagou, Vasiliki, et al.  Nature communications (2021) | 0.104 | 0.337 | 67 |
| Height | Height | 1,580,547 | Yengo, Loïc, et al.  Nature (2022) | 0.525 | -0.040 | 5049 |
| Inflammatory bowel disease | IBD | 456,414 | Wu et al.,  2021 | 0.016 | 0.003 | 26 |
| Insomnia | INS | 384,837 | Watanabe, Kyoko, et al. Nature genetics (2022) | 0.057 | 0.054 | 14 |
| Prostate cancer | PC | 140,254 | Schumacher, Fredrick R., et al.  Nature genetics (2018) | 0.143 | -0.017 | 90 |
| Parkinson’s disease | PD | 308,374 | Chang, Diana, et al.  Nature genetics (2017) | 0.025 | -0.002 | 24 |
| Rheumatoid arthritis | RA | 276,020 | Ishigaki, Kazuyoshi, et al. Nature genetics (2022) | 0.095 | 0.011 | 93 |
| Red blood cell count | RBC | 440,235 | Jiang et al.,  2019 | 0.257 | 0.091 | 759 |
| Systolic blood pressure | SBP | 427,595 | Jiang, Longda, et al. Nature genetics (2019) | 0.168 | 0.177 | 413 |
| Schizophrenia | SCZ | 130,570 | Trubetskoy, Vassily, et al., Nature (2022) | 0.438 | -0.017 | 185 |
| Type 2 diabetes | T2D | 933,970 | Mahajan, Anubha, et al. Nature genetics (2022) |  |  | 241 |
| Lung cancer | LC | 85,716 | McKay, James D., et al. Nature genetics (2017) |  |  | 14 |
| Cigarettes per day | CigDay | 325,708 | Saunders, Gretchen RB, et al. Nature (2022) |  |  | 131 |

^1^ For GWAS with non-uniform per-SNP sample sizes, we report the mean sample size across the $\sim$1 million HapMap3 SNPs analyzed.

^2^ The trait’s heritability is the genome-wide estimate from SBayesAPP obtained in the 155 cell-type analyses.

^3^ The trait’s genetic correlation with type 2 diabetes (r_g_) is the genome-wide estimate from SBayesAPP obtained in the 155 cell-type analyses.

**Supplementary Table 2.** Summary of annotation-stratified coheritability enrichment, SNP effect correlation, and shared polygenicity estimates from SBayesAPP’s analysis of type 2 diabetes (T2D) and fasting glucose, based on SNP groupings by GWAS significance.

| Annotation | Number of SNPs | coheritability enrichment | SNP effect correlation | shared polygenicity |
| --- | --- | --- | --- | --- |
| Significant | 3,993 | 702 (43) | 0.87 (0.04) | 0.21 (0.03) |
| Rest | 1,146,536 | 0.67 (0.02) | 0.46 (0.05) | 0.005 (0.0003) |
| Control | 3,993 | -0.02 (0.13) | -0.23 (0.58) | 0.01 (0.009) |

**Supplementary Table 3.** Summary of annotation-stratified coheritability enrichment, SNP effect correlation, and shared polygenicity estimates from SBayesAPP’s analysis of type 2 diabetes (T2D) and height, based on SNP groupings by GWAS significance.

| Annotation | Number of SNPs | coheritability enrichment | SNP effect correlation | shared polygenicity |
| --- | --- | --- | --- | --- |
| Significant | 29,579 | 113 (15) | -$0.23$ (0.05) | 0.26 (0.01) |
| Rest | 1,095,364 | 0.58 (0.07) | -0.09 (0.03) | 0.007 (0.0007) |
| Control | 29,579 | 0.02 (0.06) | -0.22 (0.74) | 0.0006 (0.0006) |

**Supplementary Table 4.** Estimates of polygenicity among trait-associated variants in Figure 4d.

| Trait name | Annotation name | Polygenicity among trait-associated variants (s.e.)^1^ | | | $\pi_{0}$ (s.e.) |
| --- | --- | --- | --- | --- | --- |
|  |  | T2D-specific | Other-trait-specific | Shared |  |
| T2D and fasting glucose | mature enterocyte | 0.1855 (0.0576) | 0.4814 (0.0494) | 0.3330 (0.0516) | 0.9915 (0.0047) |
|  | langerhans cell | 0.1686 (0.0503) | 0.2471 (0.0546) | 0.5843 (0.0596) | 0.9878 (0.0021) |
|  | erythrocyte | 0.1157 (0.0455) | 0.3318 (0.0688) | 0.5525 (0.0597) | 0.9692 (0.0042) |
|  | pancreatic ductal cell | 0.2574 (0.0517) | 0.3241 (0.0648) | 0.4185 (0.0358) | 0.9915 (0.0024) |
|  | pancreatic acinar cell | 0.2803 (0.0600) | 0.3099 (0.0670) | 0.4099 (0.0606) | 0.9899 (0.0013) |
|  | plasma cell | 0.2742 (0.0585) | 0.3825 (0.0526) | 0.3433 (0.0461) | 0.9832 (0.0032) |
|  | enterocyte of epithelium of large intestine | 0.2299 (0.0615) | 0.5479 (0.0789) | 0.2222 (0.0424) | 0.9922 (0.0014) |
|  | thymocyte | 0.2977 (0.0661) | 0.2657 (0.0409) | 0.4366 (0.0645) | 0.9843 (0.0037) |
|  | enterocyte of epithelium of small intestine | 0.2946 (0.0550) | 0.4298 (0.0519) | 0.2756 (0.0489) | 0.9897 (0.0014) |
|  | intestinal crypt stem cell of small intestine | 0.2398 (0.0609) | 0.4504 (0.0761) | 0.3098 (0.0644) | 0.9919 (0.0014) |
|  | pancreatic beta cell | 0.2713 (0.0674) | 0.3147 (0.0663) | 0.4139 (0.0520) | 0.9911 (0.0009) |
|  | paneth cell of epithelium of small intestine | 0.2303 (0.0660) | 0.4799 (0.0565) | 0.2899 (0.0521) | 0.9896 (0.0013) |
|  | immature natural killer cell | 0.3193 (0.0778) | 0.3175 (0.0680) | 0.3632 (0.0604) | 0.9906 (0.0030) |
|  | hepatocyte | 0.2370 (0.0540) | 0.5079 (0.0556) | 0.2552 (0.0441) | 0.9882 (0.0028) |
|  | duodenum glandular cell | 0.3060 (0.0735) | 0.4179 (0.0658) | 0.2760 (0.0397) | 0.9908 (0.0014) |
| T2D and height | mature enterocyte | 0.6369 (0.0879) | 0.2187 (0.0721) | 0.1443 (0.0403) | 0.9699 (0.0038) |
|  | langerhans cell | 0.5904 (0.1469) | 0.1322 (0.0803) | 0.2774 (0.0734) | 0.9692 (0.0144) |
|  | erythrocyte | 0.4654 (0.1501) | 0.2832 (0.1399) | 0.2514 (0.0441) | 0.9546 (0.0191) |
|  | pancreatic ductal cell | 0.7763 (0.0745) | 0.0904 (0.0336) | 0.1334 (0.0534) | 0.9689 (0.0090) |
|  | pancreatic acinar cell | 0.7206 (0.0469) | 0.1501 (0.0502) | 0.1293 (0.0247) | 0.9526 (0.0039) |
|  | plasma cell | 0.7747 (0.0497) | 0.0745 (0.0287) | 0.1508 (0.0269) | 0.9412 (0.0046) |
|  | enterocyte of epithelium of large intestine | 0.6547 (0.0630) | 0.1705 (0.0377) | 0.1748 (0.0487) | 0.9789 (0.0047) |
|  | thymocyte | 0.7347 (0.0308) | 0.1205 (0.0299) | 0.1448 (0.0227) | 0.9415 (0.0089) |
|  | enterocyte of epithelium of small intestine | 0.5995 (0.0615) | 0.2069 (0.0557) | 0.1936 (0.0333) | 0.9743 (0.0108) |
|  | intestinal crypt stem cell of small intestine | 0.6100 (0.1157) | 0.2490 (0.1094) | 0.1410 (0.0411) | 0.9692 (0.0065) |
|  | pancreatic beta cell | 0.8117 (0.0322) | 0.1116 (0.0198) | 0.0767 (0.0225) | 0.9619 (0.0038) |
|  | paneth cell of epithelium of small intestine | 0.5785 (0.0938) | 0.2462 (0.0850) | 0.1754 (0.0385) | 0.9747 (0.0032) |
|  | immature natural killer cell | 0.3742 (0.1745) | 0.2806 (0.1478) | 0.3452 (0.0473) | 0.9652 (0.0139) |
|  | hepatocyte | 0.5705 (0.0541) | 0.2126 (0.0467) | 0.2169 (0.0301) | 0.9779 (0.0019) |
|  | duodenum glandular cell | 0.6936 (0.0889) | 0.1646 (0.0738) | 0.1419 (0.0270) | 0.9526 (0.0125) |

^1^ The polygenicity among trait-associated variants are the polygenic mixture proportions scaled by $\frac{1}{1-\pi_{0}}$.

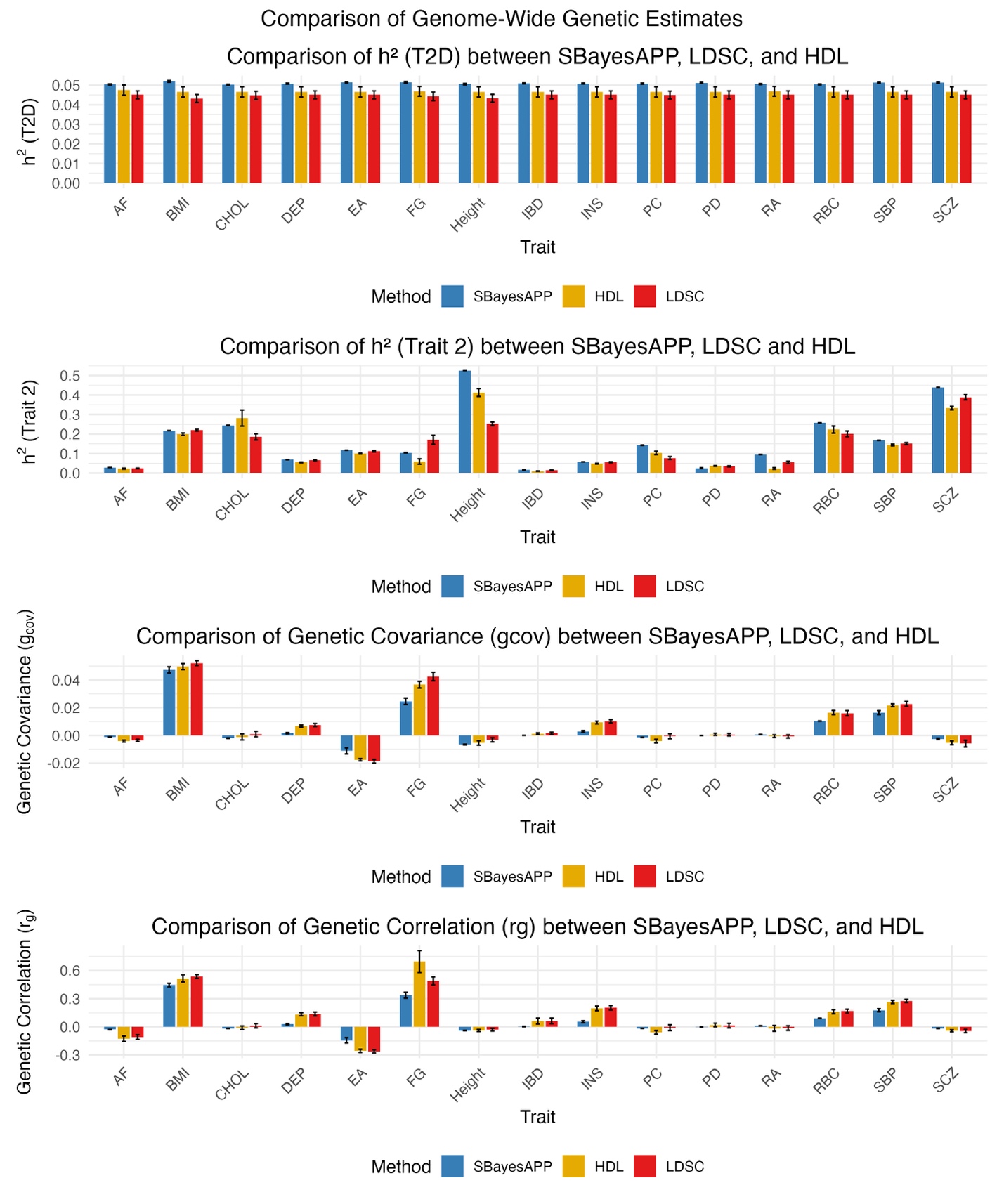

**Supplementary Figure 1.** Comparison of genome-wide heritability, genetic covariance, and genetic correlation estimates across 15 trait pairs involving type 2 diabetes (T2D), using SBayesAPP (blue), HDL (yellow), and LDSC (red). Error bars represent ±1 standard error (for HDL and LDSC) or ±1 posterior standard deviation (for SBayesAPP). The x-axis indicates the second trait in each T2D–trait pair.

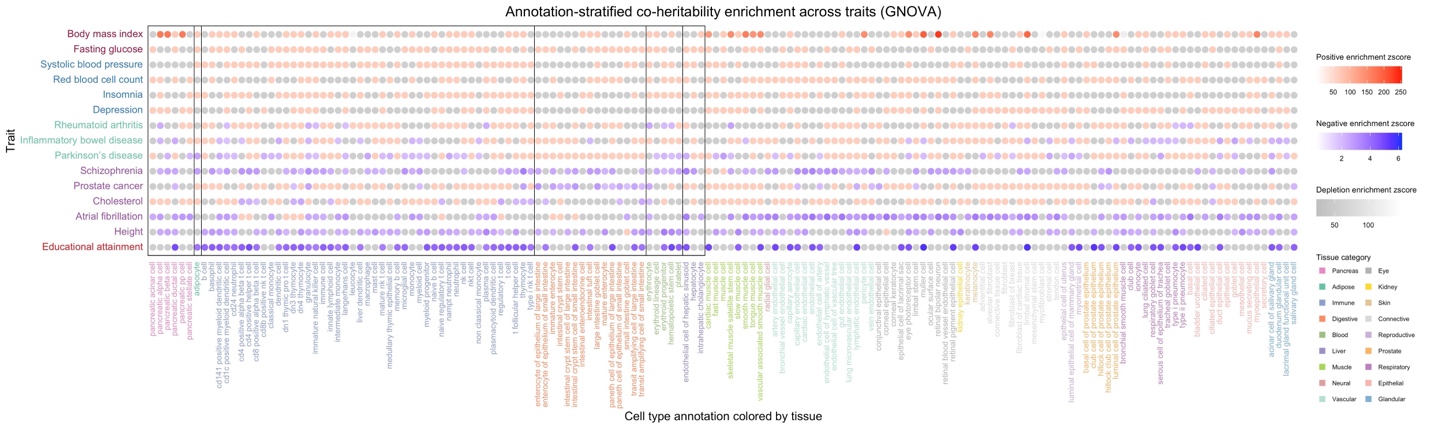

**Supplementary Figure 2.** Annotation-stratified coheritability enrichment between T2D and 15 secondary traits estimated by GNOVA. Traits are listed on the y-axis, ordered by their genome-wide genetic correlation with T2D (shown in parentheses) estimated by SBayesAPP. The x-axis displays the analyzed 155 cell type annotations grouped into 18 tissue categories. Color indicates the sign of genetic covariance and the magnitude of coheritability enrichment z-score, defined as estimates divided by standard error. Both red and blue represent enriched annotations with estimated enrichment fold greater than 1, where red corresponds to positive genetic covariance and blue to negative covariance. Darker shades of red or blue indicate higher enrichment z-scores. Gray indicates annotations with enrichment fold between –1 and 1 (i.e., depleted annotations), with darker gray representing values closer to zero (i.e., smaller z-score).

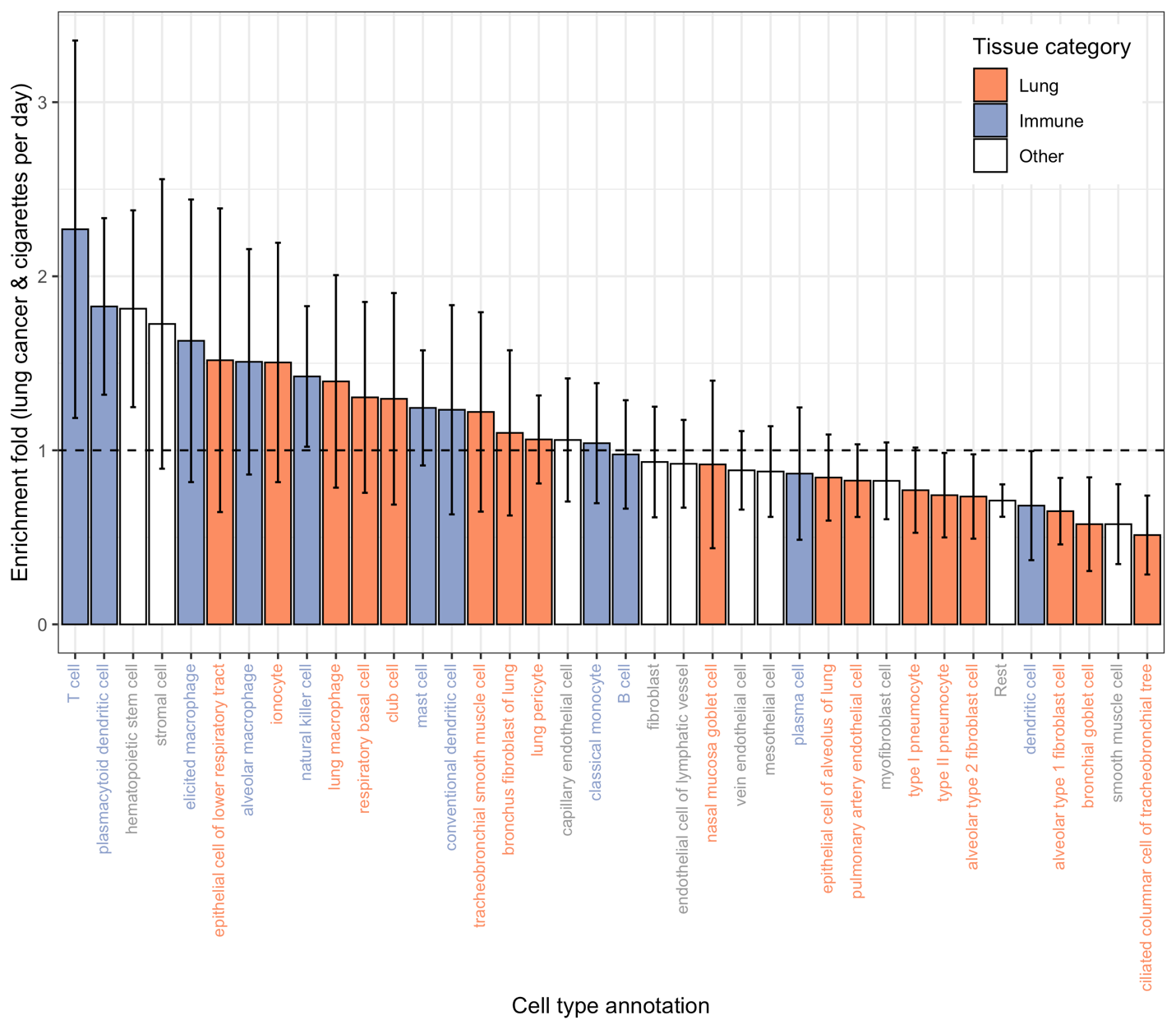

**Supplementary Figure 3.** Annotation-stratified coheritability enrichment estimates from GNOVA for the trait pair lung cancer and cigarettes per day. GNOVA estimates are shown with standard errors approximated using the delta method, based on the standard errors of corrected genetic covariance. All annotations are color-coded by tissue category.

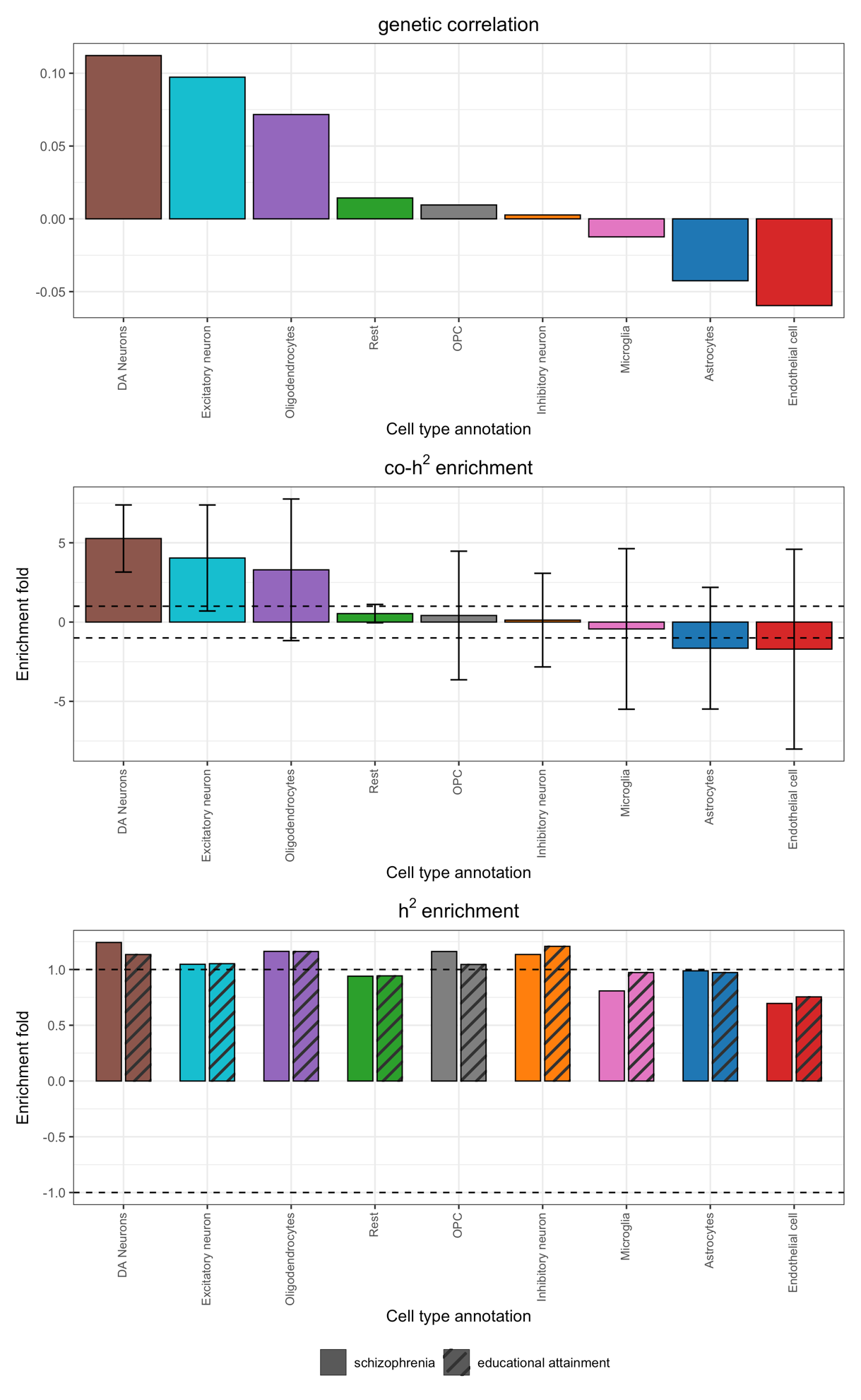

**Supplementary Figure 4**. Annotation-stratified genetic correlation, coheritability enrichment, and heritability enrichment estimated by GNOVA. GNOVA shows coheritability enrichment with standard errors obtained via the delta method from the standard errors of the corrected genetic covariance. Standard errors are not available from GNOVA for heritability or genetic correlation.

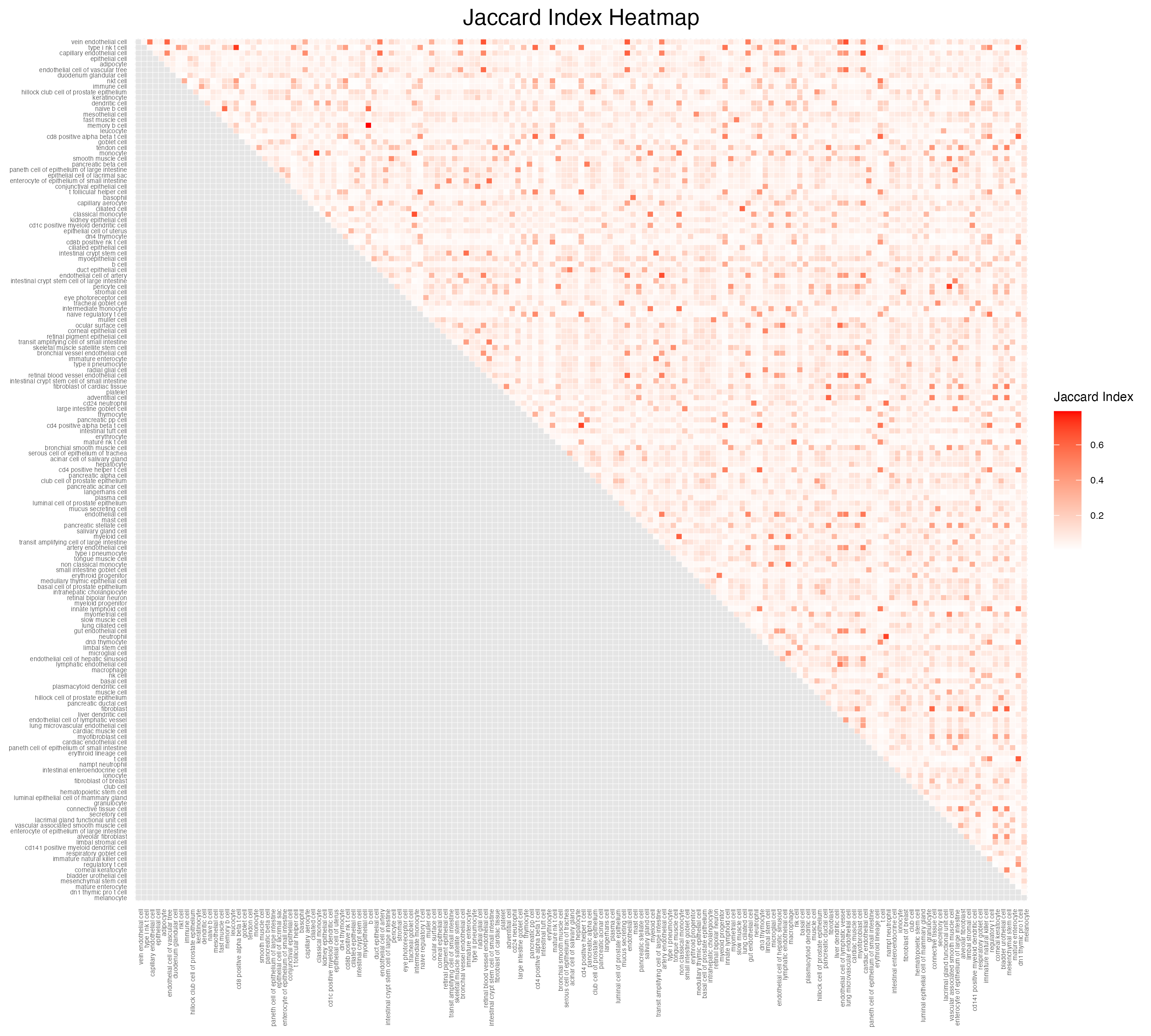

**Supplementary Figure 5.** Heatmap of the Jaccard index quantifying SNP overlap between cell-type annotation pairs. The Jaccard index is calculated as the ratio of the number of shared SNPs to the total number of unique SNPs across the two annotations.

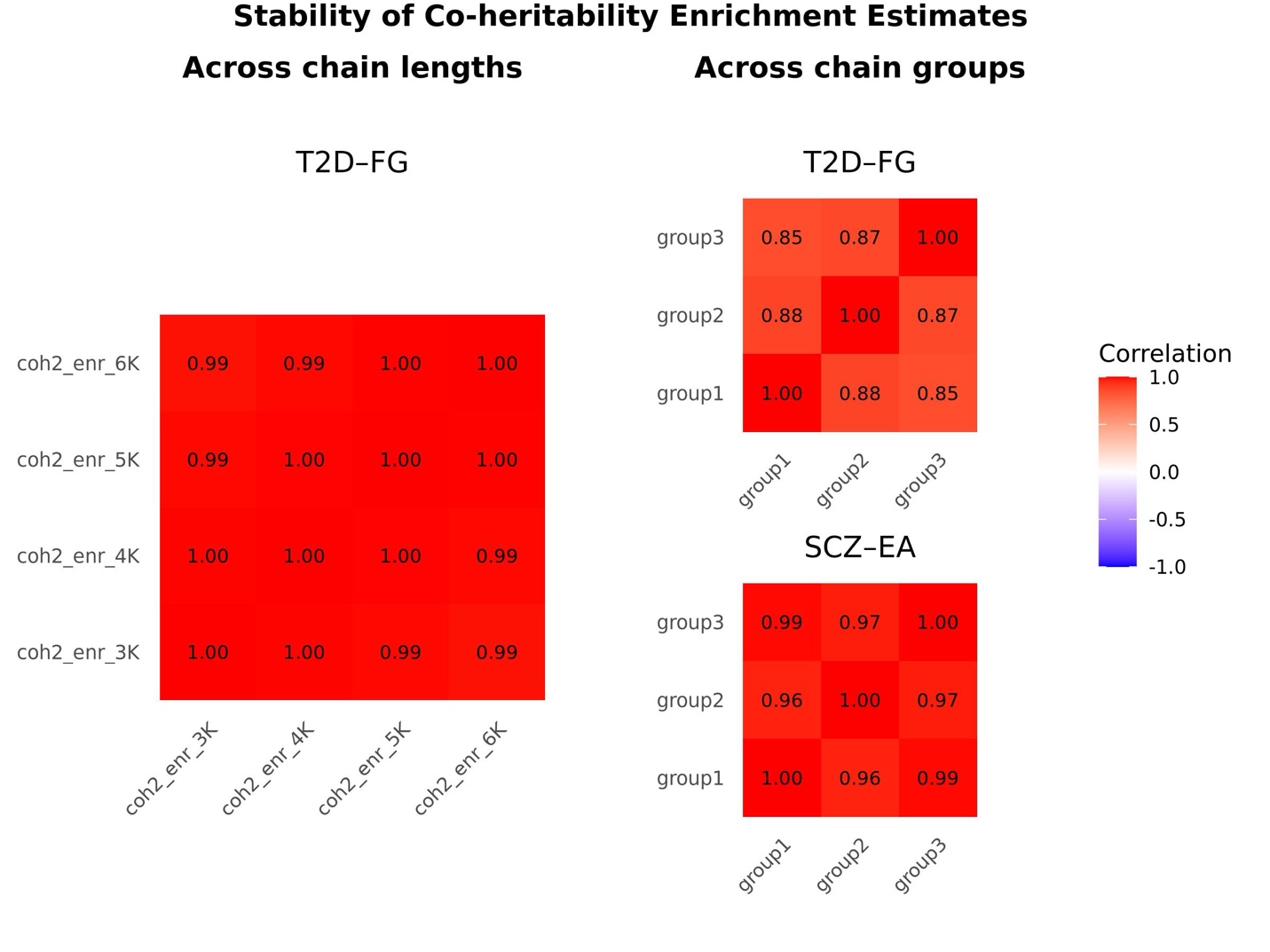

**Supplementary Figure 6. Convergence of co-heritability (co-h^2^) enrichment estimates within a chain and across grouped chains, assessed by correlation. Left**: within-chain, across lengths: Correlations of annotation-specific co-h^2^ enrichment estimates across MCMC chain lengths using the T2D–fasting glucose (FG) analysis (with 156 annotations) as a representative example. Pairwise Pearson correlations were computed among estimates from 3K, 4K, 5K, and 6K iterations within the same chain. To minimize the influence of a single dominant value, the annotation with the highest enrichment, identical across all lengths, was excluded from the correlation calculation. **Right**: Across grouped chains, T2D–FG and SCZ–EA: Guided by left panel, which supports 3,000 iterations as sufficient for stable single-chain estimation in real data, we computed correlations across groups of independent chains. The heatmaps show pairwise Pearson correlations of co-heritability enrichment across three groups of MCMC runs, each group comprising 10 independent chains of 3,000 iterations, for T2D–FG analysis (with 156 annotations) and schizophrenia–educational attainment analysis (with 9 annotations).
